# PCMD-1 stabilizes the PCM scaffold and facilitates centriole separation

**DOI:** 10.1101/2024.11.14.623570

**Authors:** Alina Schreiner, Astrid Heim, Friederike Wolff, Esther Zanin, Tamara Mikeladze-Dvali

## Abstract

Centrosomes are highly dynamic organelles, and maintaining their stability is crucial for spindle pole integrity and bipolar spindle formation. Centrosomes consist of a pair of centrioles surrounded by the pericentriolar material (PCM). In *Caenorhabditis elegans*, interactions between the PCM scaffold protein SPD-5 and the regulatory kinase PLK-1 are essential for PCM formation and disassembly. However, how PCM stability is established and maintained remains an open question. Here, we address this question by analyzing the function of PCMD-1, a protein that predominantly localizes to centrioles. Mutations in the predicted PLK-1 binding sites of PCMD-1 result in a reduction of centrosomal PLK-1 and PCMD-1 levels, leading to severe distortion of the PCM scaffold. The disorganization of the PCM is already evident during its formation and results in the assembly of a structurally unstable mitotic centrosome, which is unable to resist the microtubule pulling forces exerted on the spindle pole. As a consequence of a weakened PCM scaffold, the pulling forces are not effectively relayed to the entrapped centriole pair, resulting in delayed centriole separation in late anaphase. Together, these findings show that PCMD-1, which predominantly localizes to centrioles and tethers the PCM scaffold to them, is essential for stabilizing the entire PCM scaffold and ensuring timely centriole separation during PCM disassembly. We propose a model in which PCMD-1 initiates the ordered assembly of the PCM scaffold by biasing the PCM core to a certain intrinsic order around the centrioles. This intrinsic order acts as a seed that propagates throughout the surrounding micron-scale PCM scaffold, providing the necessary strength and structural integrity for the centrosome.

## INTRODUCTION

Centrosomes are the main microtubule organizing centers of animal cells that orchestrate the assembly of a bipolar spindle during cell division. Centrosomes comprise a pair of centrioles, surrounded by the pericentrosomal material (PCM). While the PCM nucleates and tethers spindle microtubules, it must also withstand extremely high pulling forces exerted on the centrosome by the emanating microtubules (Grill et al., 2001; Laan et al., 2012). Maintaining a stable, integer centrosome is pivotal for coordinated cell division and faithful chromosome segregation. Several different mechanisms can cause the formation of a spindle apparatus with multiple poles, which is a source of aneuploidy and has been often observed in cancer cells (Ganem et al., 2009; Holland and Cleveland, 2009). Multipolar spindles can be the result of numerical aberrations or premature separation of centrioles, or alternatively be caused by the fragmentation of an unstable and weakened PCM, leading to the assembly of extra, acentriolar poles (Maiato and Logarinho, 2014). How PCM stability is achieved, such that it can withstand the microtubule pulling forces without fragmentation, is still not fully understood. The PCM is a proteinaceous matrix, undergoing dynamic changes during the cell cycle. The interphase PCM forms a highly structured core around the centrioles (Fu and Glover, 2012; Lawo et al., 2012; Mennella et al., 2012; Sonnen et al., 2012). Upon mitotic entry, the PCM undergoes a maturation process and expands to its maximum extent at metaphase. With the entry into anaphase the PCM fractures and dissolves leaving behind a small PCM core surrounding the centriole.

Posttranslational modifications of PCM components drive this dynamic process and are intrinsically intervened with PCM stabilization during mitosis. Aurora A and PLK1 (Polo-like Kinase) are two multifunctional mitotic kinases linked to centrosome integrity. In human U2OS cells, the decrease of Aurora A activity and its substrate ch-TOG leads to the loss of centrosome stability and the formation of extra spindle poles (De Luca et al., 2008; Foraker et al., 2012; Ou et al., 2018). Along with its diverse roles in cell division, such as the promotion of mitotic entry, nuclear envelope breakdown (NEBD), sister chromatid separation, and cytokinesis, PLK1 promotes centrosome maturation and centriole separation (Joukov and De Nicolo, 2018). Increased PCM fragmentation has been observed in PLK1-inhibited cultured and zebrafish cells (Aljiboury et al., 2022; Rathbun et al., 2020). A possible substrate for PLK1 in terms of centrosome stability is the protein Kizuna (Kiz) (Oshimori et al., 2006; Thomas et al., 2014). Kiz tethers the PCM to the centrioles in human cells and its phosphorylation by PLK1 is needed to maintain spindle pole integrity. Another key substrate of PLK1 is human pericentrin (PCNT) (Barrera et al., 2010; Lee and Rhee, 2012; Pagan et al., 2015). Phosphorylation of PCNT by PLK1 is a prerequisite for its separase-based cleavage towards the end of mitosis. This leads to timely PCM disassembly and subsequent centriole disengagement and separation (Kim et al., 2015; Lee and Rhee, 2012; Matsuo et al., 2012; Tsou et al., 2009). In agreement with this, in HeLa cells depleted of PCNT, centrioles separate prematurely (Kim et al., 2019). Premature centriole separation was also observed in Pericentrin-Like-Protein (PLP) mutant sensory organ precursors of flies (Roque et al., 2018). Interestingly, PLP mutant embryos recruit PCM to the centrosomes but exhibit a highly fragmented and flared PCM, indicating that proper PLP levels are not only necessary for timely centriole separation but also to maintain centrosome stability (Richens et al., 2015).

Research in *C. elegans* led to the proposal that the centrosome can be best described as a viscoelastic entity, with physical properties reminiscent to a non-Newtonian fluid (Mittasch et al., 2020; Woodruff, 2021). According to the current model, after maturation and growth, the PCM transitions from a strong and more ductile physical state at metaphase to a weak and deformable one in anaphase. The physical properties at metaphase allow the PCM to withstand deformation by microtubule pulling forces. The same forces can easily disassemble the brittle PCM in anaphase and release the entrapped pair of centrioles (Magescas et al., 2019).

Spindle-Defective-5 (SPD-5), the homolog of Cdk5Rap2 in humans and Cnn in *Drosophila* and PLK-1 are major drivers of PCM dynamics in *C. elegans.* SPD-5 is the main scaffold protein of the PCM matrix that can self-assemble and form micron-scale networks *in vitro* (Hamill et al., 2002; Woodruff et al., 2015). Network formation of purified SPD-5 is highly accelerated by the presence of the structural protein SPD-2 and by PLK-1 phosphorylation. SPD-5 contains a phospho-regulated-multimerization (PReM) domain and a C-terminal centriole targeting sequence. The PReM domain mediates the interaction of SPD-5 with PLK-1 and phosphorylation by PLK-1 finetunes the complex landscape of SPD-5 macromolecular self-interactions *in vitro* (Nakajo et al., 2022; Rios et al., 2024; Woodruff et al., 2015).

*In vivo* SPD-2, along with the kinases PLK-1 and AIR-1 (Aurora A), are required for the centrosome scaffold assembly and maturation (Cabral et al., 2019; Hannak et al., 2001; Kemp et al., 2004; Özlü et al., 2005; Pelletier et al., 2004). PLK-1 is recruited to the centrioles by SPD-2 (Decker et al., 2011). Phosphorylation of SPD-5 by PLK-1 at four residues drives PCM expansion at mitotic entry (Woodruff et al., 2015; Wueseke et al., 2016). When these residues are rendered phosphodeficient, SPD-5 still seeds the PCM core around the centrioles, but is unable to assemble an expanded micron-scale scaffold. PLK-1, in contrast to AIR-1 and SPD-2, is not only needed for centrosome maturation but also to maintain an already assembled PCM (Cabral et al., 2019). Acute inhibition of PLK-1 at metaphase causes rapid PCM dispersal and disassembly. Centrosomal PLK-1 is downregulated at anaphase in a CDK-1- dependent manner (Cabral et al., 2019), and concomitant with this, the PCM weakens and starts to disassemble. The switch between PLK-1 activity at the centrosome and its counteracting phosphatase PP2A is thought to trigger PCM fragmentation and eventual dissolution by the microtubule pulling forces (Enos et al., 2018; Magescas et al., 2019; Mittasch et al., 2020). Interestingly, when centrioles are ablated at early time points of centrosome maturation, the remaining PCM cannot recover, it disperses prematurely even before anaphase onset. The dispersal can be reversed by microtubule depolymerization, indicating that a fragile and untethered PCM scaffold dissipates in a microtubule-dependent manner, however, still retains its intrinsic ability to form a spherical entity (Cabral et al., 2019).

While much progress has been made in understanding the role of the scaffold protein SPD-5 and polo kinase PLK-1 in regulating dynamical changes in mechanical properties of the PCM, how other centrosomal proteins contribute to this process and how a weakened PCM affects centriole separation is less well understood.

The PCM scaffold is tethered to the centrioles by the intrinsically disordered protein Pericentriolar matrix deficient-1 (PCMD-1) (Erpf et al., 2019; Stenzel et al., 2021). One-cell embryos mutant for *pcmd-1* are 100% embryonic lethal, due to the failure of SPD-5 recruitment to the centrioles, absence of the PCM core formation, and failure of PCM scaffold self-assembly. Occasionally SPD-5 forms an extremely compromised PCM, impeding an analysis of PCMD-1 function in maintaining PCM integrity. We have previously shown that PCMD-1 binds SPD-5 and PLK-1 and is also able to recruit SPD-5 and PLK-1 to an ectopic cellular location (Erpf et al., 2019; Stenzel et al., 2021). Centrosomal PLK-1 levels are also decreased in *pcmd-1(t3421)* mutant embryos, suggesting that along with SPD-2, PCMD-1 contributes to PLK-1 recruitment to the centrosome and thereby facilitates its centrosome-related functions (Erpf et al., 2019; Stenzel et al., 2021).

Here we show that the interaction of PCMD-1 with PLK-1 is necessary to establish a coherent and stable PCM *in vivo*. When predicted PLK-1 binding sites are mutated in PCMD-1, PLK-1 levels at the centrosome are not maintained. The PCM core is seeded and the scaffold is recruited to centrioles but assembles into a fragile and distorted network that cannot withstand microtubule pulling forces and is easily deformed by exaggerated forces applied to spindle poles. As a consequence of a weakened PCM scaffold, pulling forces are not relayed to the entrapped centriole pair, resulting in a delayed centriole separation in anaphase, suggesting a key role of PCMD-1 in centriole separation.

## RESULTS

### Mutating putative PLK-1-binding sites in PCMD-1 leads to severe PCM scaffold disorganization

Downregulating PLK-1 function has pleiotropic effects on the embryo and does not allow us to test the functional importance of the PLK-1-PCMD-1 interaction. To overcome this, we set out to identify and mutate predicted PLK-1 Polo Box Domain (PBD) binding sites in PCMD-1. PLK-1 PBD-sites are phosphorylation sites, usually primed by PLK-1 itself or another kinase (Garcia-Alvarez et al., 2007). Using a prediction algorithm and manual annotation, we identified 23 potential PBD consensus sites in PCMD-1 and rendered all sites phosphodeficient, by changing them to alanines (Table 1). This phosphodeficient transgene will be referred to as PCMD-1- PBD^m^ (for PBD mutant) throughout the manuscript (Figure 1 A). First, we tested the modified protein by transiently expressing it in HEK293T cells. EGFP-tagged PCMD- 1^WT^ localized to microtubules (Figure S1 A) and was able to recruit mCherry::PLK-1 and mCherry::SPD-5 to this ectopic location in 92% and 100% of the cells, respectively. Interestingly, EGFP::PCMD-1-PBD^m^ localized to microtubules with the same efficiency, however, it was less efficient in recruiting its binding partners (Figure S1 A). mCherry::PLK-1 localized to the ectopic location in only 23% of the cells and mCherry::SPD-5 never localized to EGFP::PCMD-1-PBD^m^ decorated microtubules but instead formed aggregates in the cytoplasm. Encouraged by these findings, we used the previously established ‘single-copy-replacement-system’ to assess the functionality of the modified protein in *C. elegans* (Stenzel et al., 2021). In short, N-terminally GFP-tagged PCMD-1-PBD^m^ was crossed into the *pcmd-1(syb975)* null mutant background and the wild type version (GFP::PCMD-1^WT^) expressed in the same mutant background was used as a control.

**Figure 1:**
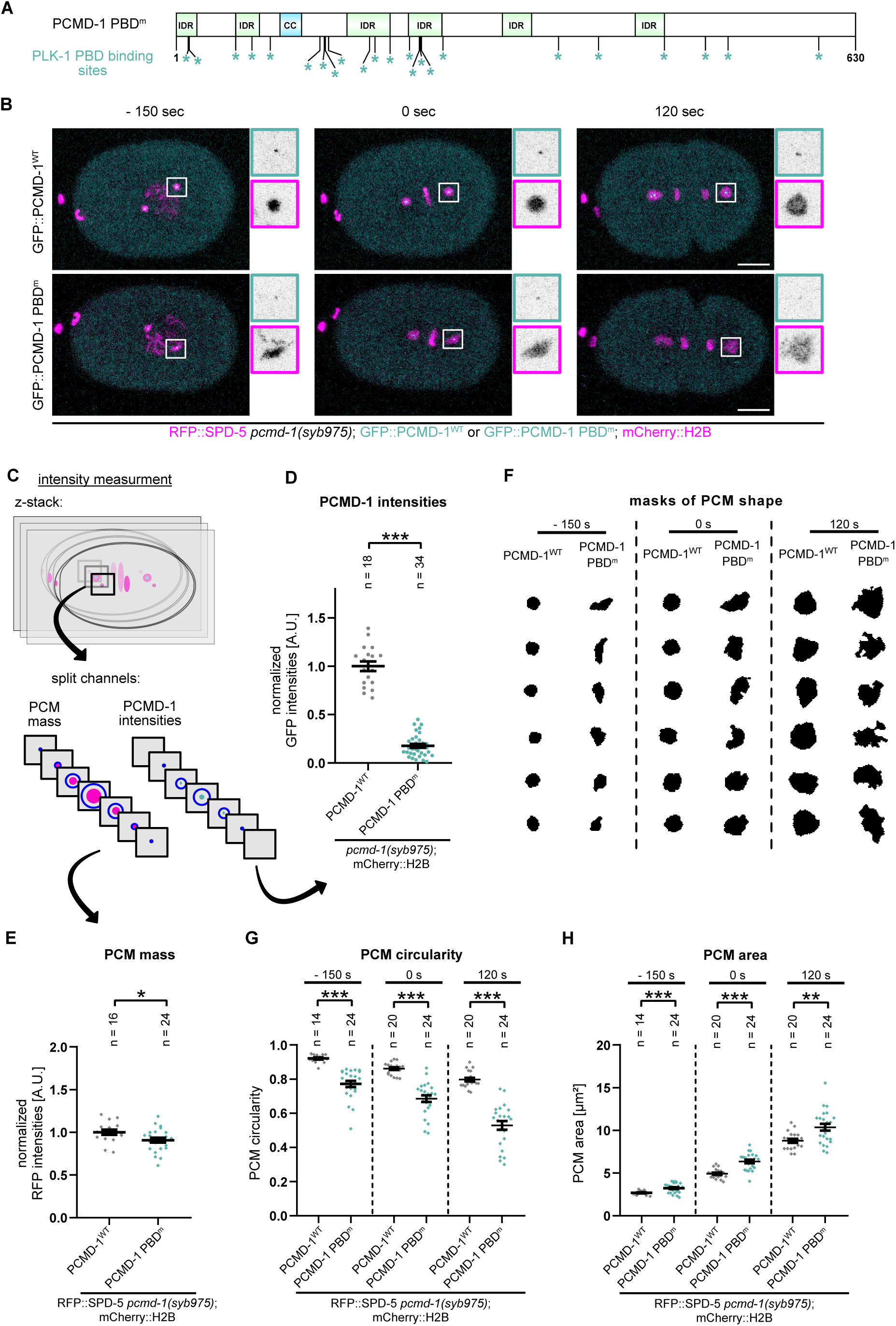
PCMD-1-PBD^m^ leads to severe PCM scaffold disorganization. A) Schematic representation of PCMD-1^WT^ structure, with predicted intrinsically disordered regions (IDR) in green and the coiled-coil region (CC) in blue. The position of the mutated PBD^m^ binding sites is indicated by an asterisk under the protein structure. B) Stills of timelapse movies of embryos expressing GFP::PCMD-1^WT^ and GFP::PCMD-1 PBD^m^ in combination with mCherry::H2B and RFP::SPD-5 in the *pcmd-1(syb975)* background. Centrosomal areas are shown enlarged in insets. Scale bar = 10 μm. C) Schematics of quantifications of PCM mass and PCMD-1 intensities. D) Normalized GFP intensity values of GFP::PCMD-1^WT^ and GFP::PCMD-1 PBD^m^ at metaphase. p-value from a Wilcoxon rank sum test. E) Normalized PCM mass at the centrosome during metaphase. p-value from Two Sample t-test. F) Representative examples of PCM shapes at indicated timepoints. G) Quantification of PCM circularities at indicated time points. p-values from Wilcoxon rank sum testand Welch Two Sample t-test. H) Quantification of PCM area at indicated time points. p-values from Two Sample t-test or Welch Two Sample t-test. For all movies: 0 sec is metaphase. n = number of analyzed centrosomes.

We assessed the functionality of GFP::PCMD-1-PBD^m^ by scoring the rescue of embryonic viability. Compared to wild type worms, none of the embryos laid by *pcmd-1(syb975)* mutants worms hatch at restricted temperature (Figure S1 B). Expressing GFP::PCMD-1^WT^ in this background rescued embryonic viability to 97.7%. In comparison only 86.4% of GFP::PCMD-1-PBD^m^ embryos were viable (Figure S1 B). This result indicates that by mutating PLK-1-PBD-sites the protein lost some of its functionality. Then, we analyzed the recruitment of the GFP::PCMD-1-PBD^m^ by measuring relative GFP intensities at the centrosome (Figure 1 B, C and D). GFP::PCMD-1-PBD^m^ centrosomal levels were 5,5 times reduced compared to the GFP::PCMD-1^WT^, even though the transgenes were expressed at comparable levels (Figure S1 C).

Most of the *pcmd-1* mutant embryos fail to recruit the PCM core to the centrioles prior to mitotic entry (Erpf et al., 2019; Stenzel et al., 2021). To test whether the reduced lethality of PCMD-1-PBD^m^ transgenes could be explained by a failed or compromised PCM recruitment of the *pcmd-1(syb975)* mutants, we crossed RFP::SPD-5 into the strains and performed live-cell imaging during the first cell cycle of the embryos (Figure 1 B, Videos 1 and 2). Similar to the control, all GFP::PCMD-1-PBD^m^ embryos were able to recruit SPD-5 to the centrioles (n=20 PBD^WT^, n=22 PBD^m^). The PCM mass at metaphase was only slightly decreased, indicating that in contrast to the HEK293T cells, almost wildtype levels of RFP::SPD-5 PCM scaffold are recruited to centrosome in the *C. elegans* embryo (Figure 1 B, C and E). Therefore, mutating predicted PBD-sites did not significantly interfere with the PCM tethering function of PCMD-1. However, we noticed that the PCM scaffold recruited by GFP::PCMD-1-PBD^m^ was severely disorganized, showing flared extensions. To better characterize the distortion of PCM scaffold, we quantified the PCM shape parameters, such as circularity and area, at prometaphase (-150 sec), metaphase (0 sec) and late anaphase (120 sec) by creating masks of RFP::SPD-5 (Figure 1 C and F). In control embryos the centrosome is almost circular with a circulatory value close to 1 in prometaphase and metaphase. Mean circularity slightly decreases from 0.9 in metaphase to 0.8 when centrosomes enter anaphase (Figure 1 F and G). In GFP::PCMD-1-PBD^m^ embryos the PCM scaffold fails to assemble into a spherical structure from the beginning on. Mean circularity of 0.8 at prometaphase is comparable to the mean circularity of control centrosomes in anaphase, and it decreases even more dramatically when embryos progress to anaphase (Figure 1 F and G). Accordingly, the area of the PCM scaffold, is significantly larger than the control at every timepoint analyzed (Figure 1 F and H).

In summary, this indicates that embryos expressing PCMD-1-PBD^m^ are viable, albeit the viability is lower than in control embryos. PCMD-1-PBD^m^ is less efficiently recruited to the centrioles, however these levels are sufficient to tether the PCM scaffold. The recruited PCM appears distorted and severely disorganized. Together, by mutating predicted PBD-sites of PLK-1 in PCMD-1 we have created a setting where the PCM scaffold is recruited to centrioles at close to normal levels, and therefore it allows us, for the first time, to analyze the function of PCMD-1 in scaffold formation.

### PLK-1 is not maintained at the PCMD-1-PBD^m^ centrosomes

We showed previously that PLK-1 levels at the centrosome are reduced by 60%, but not entirely eliminated when PCMD-1 function is compromised. The remaining PLK-1 in *pcmd-1(t3421)* mutant embryos is recruited by SPD-2 (Erpf et al., 2019). Different to PCMD-1^WT^, PCMD-1-PBD^m^ is unable to recruit PLK-1 to an ectopic location, when expressed in HEK293T cells (Figure S1 A). To explore whether by mutating PBD-sites in PCMD-1, recruitment and maintenance of PLK-1 at the centrosome is restored, we made use of a strain expressing an endogenously mCherry-tagged PLK-1.

In PCMD-1^WT^ embryos, PLK-1::mCherry is recruited during prophase and reaches its maximal levels at metaphase (0 sec) (Figures 2 A). Although cytoplasmic PLK-1 levels are higher at the anterior part of the embryo (Budirahardja and Gonczy, 2008; Rivers et al., 2008), we did not detect any difference between PLK-1 levels at the anterior versus posterior centrosome at metaphase (Figures 2 A and 2B, note distribution of non-circled and circled dots, respectively). In late anaphase (120 sec) PLK-1 levels at both centrosomes dropped and were undetectable (Figures 2 A). In PCMD-1-PBD^m^ embryos PLK-1 was recruited to the centrosome in prometaphase, however its levels were significantly reduced by nearly 30% at metaphase (Figures 2 A and 2 B). These data indicates that intact PCMD-1 is needed to maintain wild type PLK-1 levels specifically at the metaphase centrosome. Since PLK-1 played a key role in nuclear envelope break down, altered PLK-1 levels could impact cell cycle entry and progression (Martino et al., 2017). To rule out this possibility, we measured the length of the first two cell cycles of the PCMD-1^WT^ and PCMD-1-PBD^m^ embryos, and found no significant difference in cell cycle progression between the two strains (Figure S1 D).

**Figure 2:**
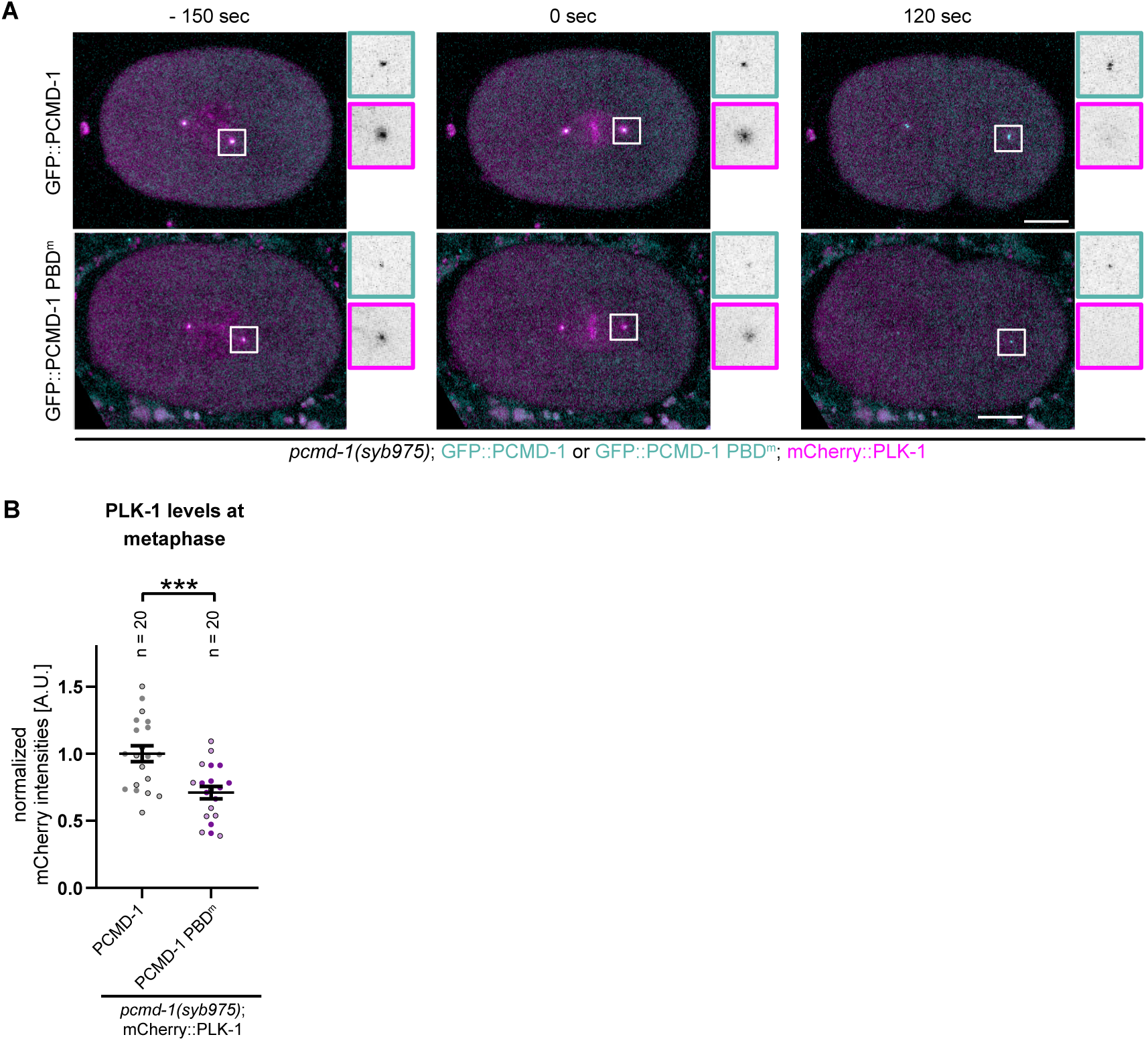
PLK-1 is not maintained at the centrosomes in PCMD-1-PBD^m^ embryos. A) Stills of timelapse movies of embryos expressing GFP::PCMD-1^WT^ and GFP::PCMD-1 PBD^m^ in combination with PLK-1::mCherry in the *pcmd-1(syb975)* background. Centrosomal areas are shown enlarged in insets. Scale bar = 10 μm. B) Normalized mCherry::PLK-1 intensities at the centrosome at metaphase. Circled points indicate posterior centrosomes. p-value from Two Sample t-test. For all movies: 0 sec is metaphase. n = number of analyzed centrosomes.

### Phosphorylation of PCMD-1 by PLK-1 is not required for PCM scaffold integrity

Since PCMD-1 binds PLK-1, PLK-1 could in turn phosphorylate PCMD-1 and this phosphorylation could contribute to PCM stabilization. We first tested whether, PLK-1 is at all able to phosphorylate PCMD-1. To do so we performed an *in vitro* kinase assay (Figure S2 C). Shortly, we incubated purified PCMD-1 with active PLK-1 kinase and γ^32P^-ATP. Radiographs clearly indicate a strong phosphorylation signal at the molecular weight of PCMD-1. Therefore, PLK-1 not only binds, but also phosphorylates PCMD-1 *in vitro*. To identify PLK-1 phosphorylation sites in PCMD-1, we used a prediction program and manual annotation. Using a medium threshold, we identified 30 putative sites and, rendered them phosphorylation deficient by mutating all the sites to alanins (Table 1). As for PCMD-1-PBD^m^, the GFP-fused transgene was expressed in the *pcmd-1(syb975)* null background using the ‘single-copy-replacement system’ (Figure S2 A and B, Video 3). We refer to this transgene as PCMD-1-PD^m^, for phosphorylation deficient.

To assess its functionality, we assayed PCMD-1 levels and the rescue of the *pcmd-1(syb975)* lethality compared to the control strain. Interestingly, neither PCMD-1 centrosomal level, nor embryonic survival rates or cell cycle progression differed from the control (Figures S2 D and E, S1 D). Second, we assayed PCM mass and shape parameters (Figures S2 F, G and H). At all timepoints analyzed PCM mass, as well as PCM circularity and area were comparable to the control embryos.

We conclude that mutating nearly 5% of the disordered protein and rendering the predicted PLK-1 phosphorylation sites phosphodeficient, neither impacts the functionality of PCMD-1, nor changes the PCM scaffold integrity and therefore we focused on the function of PLK-1 binding with PCMD-1.

### Distorted PCM scaffold responds to the modulation of microtubule pulling forces

A labile PCM scaffold could fail to resist to forces excreted by the microtubules and this could explain its disorganization and flaring at different mitotic stages. To test this hypothesis, we decided to alter the pulling forces exerted by the microtubules.

To decrease microtubule pulling forces, we used RNAi treatment against the G protein activators GPR-1/2 (Colombo et al., 2003; Gotta et al., 2003; Grill et al., 2003; Tsou et al., 2003), and mock RNAi was used as a negative control (Figures 3 A and B). If microtubule pulling forces are responsible for the PCM scaffold distortion, we expect that the circularity and PCM area will be restored when these forces are reduced. The effectiveness of *gpr-1/2* RNAi was confirmed by the failed posterior spindle displacement at metaphase (Panbianco et al., 2008) (Figure S3 A). Interestingly, at prometaphase and metaphase, the mean circularity of the PCM in *gpr-1/2* RNAi-treated PCMD-1-PBD^m^ embryos were not significantly different from the mock-treated ones, meaning that the diminished microtubule pulling forces are unable to restore circularity (Figures 3 C and D). Mean circularity at both stages had a lower value than the one of the PCMD-1^WT^ embryos. In anaphase, in PCMD-1^WT^ embryos treated with *gpr-1/2* RNAi, circularity was restored to the values of metaphase. Similarly, mean circularity in *gpr-1/2* RNAi-treated PCMD-1-PBD^m^ embryos was restored, albeit to a value comparable with a disassembling PCM in mock-treated PCMD-1^WT^ embryos. A similar trend was observed with the PCM area (Figure 3 C and E). No significant changes were observed in PCMD-1-PBD^m^ embryos with *gpr-1/2 (RNAi)* in the prometaphase and metaphase. In late anaphase the mean values of the *gpr-1/2* RNAi-treated PCMD-1-PBD^m^ embryos were restored and fell slightly under the ones of mock-treated PCMD-1^WT^.

**Figure 3:**
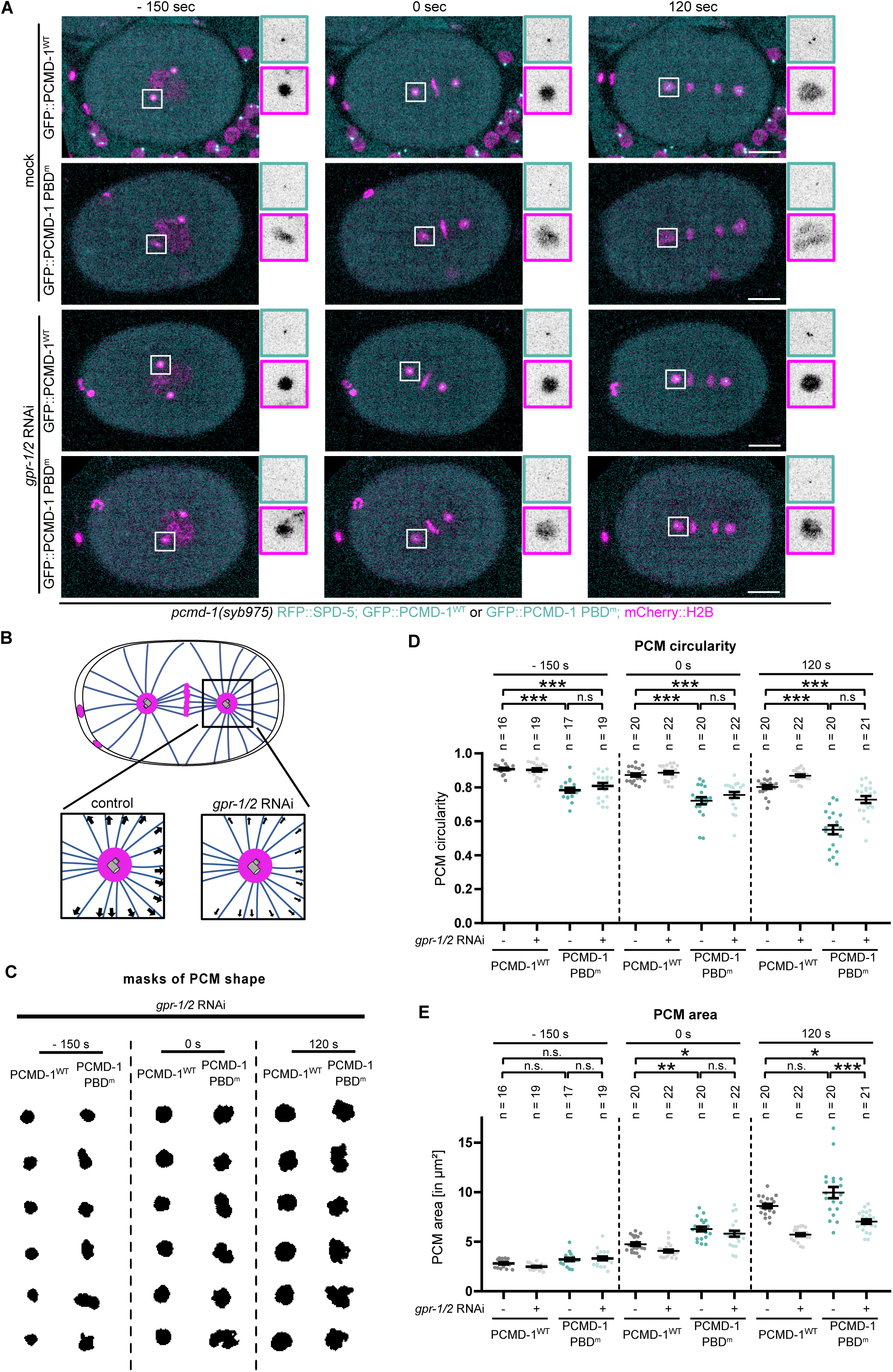
Decreased microtubule pulling forces partially rescue PCM scaffold distortion of PCMD-1-PBD^m^ centrosomes. A) Stills of timelapse movies of embryos expressing GFP::PCMD-1^WT^ and GFP::PCMD-1 PBD^m^ in combination with mCherry::H2B and RFP::SPD-5 in the *pcmd-1(syb975)* background and treated with mock or *gpr-1/2* RNAi. Centrosomal areas are shown enlarged in insets. Scale bar = 10 μm. B) Schematics of change in microtubule pulling forces in embryos treated with mock or *gpr-1/2* RNAi. C) Representative examples of PCM shapes under *gpr-1/2* RNAi at indicated timepoints. D) Quantification of PCM circularities at indicated time points. p-values from Kruskal Wallis test with Dunn post-hoc. E) Quantification of PCM area at indicated time points. p-values from Kruskal Wallis test with Dunn post-hoc. For all movies: 0 sec is metaphase. n = number of analyzed centrosomes.

A labile PCM scaffold should be easier deformable in the presence of increased microtubule-pulling forces. To increase microtubule-pulling forces we knocked down the casein kinase 1 gamma (CSNK-1), a kinase negatively regulating GPR-1/2 localization to the cortex. In *csnk-1* RNAi cortical levels of GPR-1/2 are elevated and equalized, leading to an increase of pulling forces exerted at both spindle poles (Panbianco et al., 2008) (Figures 4 A and B). The effectiveness of *csnk-1* RNAi treatment was confirmed by monitoring aberrant spindle displacement, excessive spindle rocking and flattening of the anterior spindle pole (data not shown).

**Figure 4:**
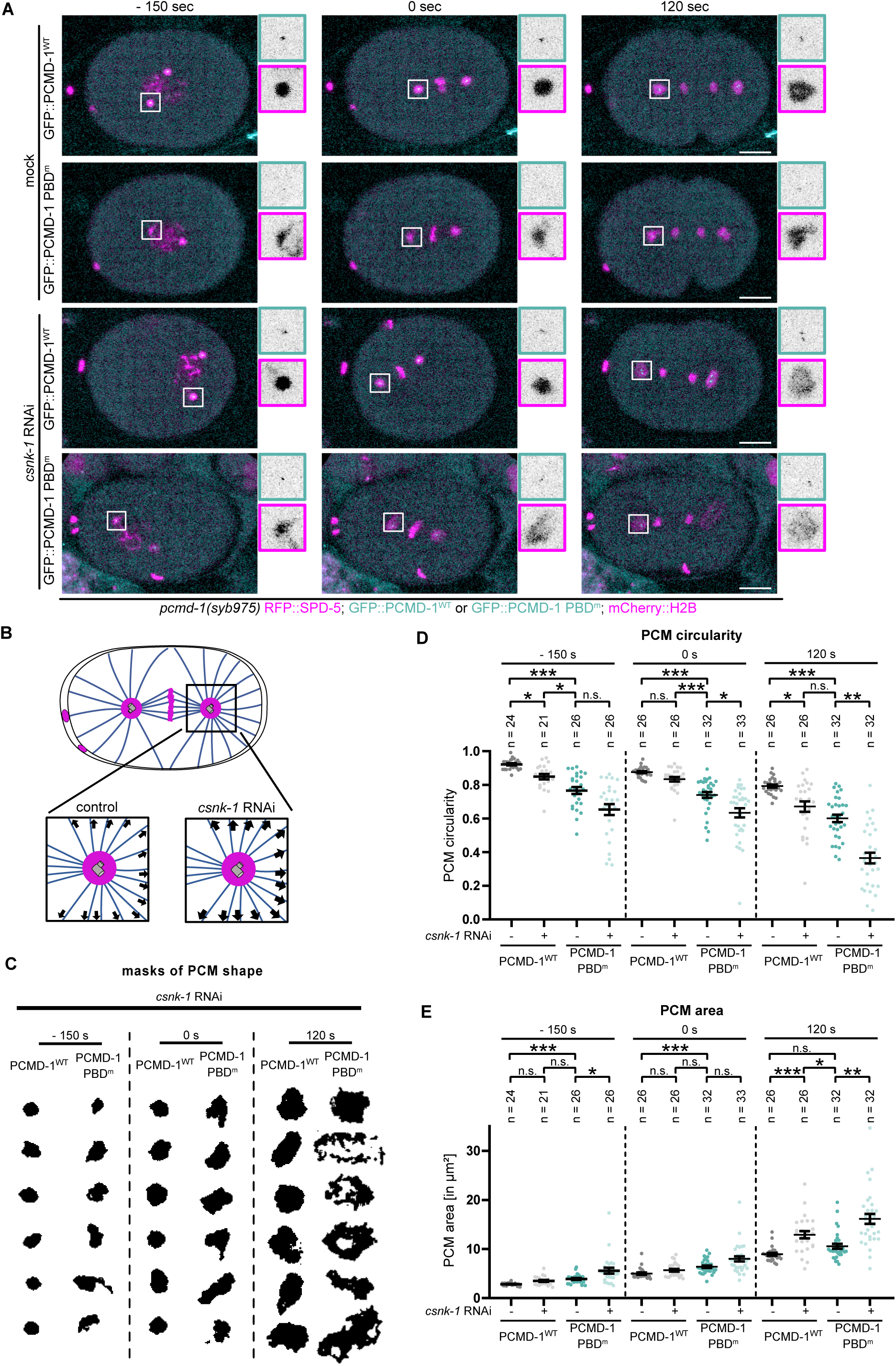
PCMD-1-PBD^m^ centrosomes cannot counteract increased microtubule pulling forces. A) Stills of timelapse movies of embryos expressing GFP::PCMD-1^WT^ and GFP::PCMD-1 PBD^m^ in combination with mCherry::H2B and RFP::SPD-5 in the *pcmd-1(syb975)* background and treated with mock or *csnk-1* RNAi. Centrosomal areas are shown enlarged in insets. Scale bar = 10 μm. B) Schematics of change in microtubule pulling forces in embryos treated with mock or *csnk-1* RNAi. C) Representative examples of PCM shapes under *csnk-1* RNAi at indicated timepoints. D) Quantification of PCM circularities at indicated time points. p-values from Kruskal Wallis test with Dunn post-hoc. E) Quantification of PCM area at indicated time points. p-values from Kruskal Wallis test with Dunn post-hoc. For all movies: 0 sec is metaphase. n = number of analyzed centrosomes.

In PCMD-1^WT^ embryos the PCM scaffold progressively deforms and shows decreased mean circularity and increased area values when higher pulling forces are applied (Figures 4 D and E). Interestingly, circularity values of *csnk-1(RNAi)*-treated PCMD-1^WT^ centrosomes at prometaphase and metaphase reveal less distortion compared to mock RNAi-treated PCMD-1-PBD^m^ centrosomes. This indicates that even increased microtubule pulling forces are unable to distort the wild type PCM to the same degree as PCMD-1-PBD^m^ centrosome.

When microtubule pulling forces were increased in PCMD-1-PBD^m^ embryos, the already compromised PCM deformed even more dramatically (Figures 4 A, C, D and E). From prometaphase going into metaphase, an extremely high degree of variability in mean circularity and area became evident. However, the impact was most obvious in anaphase, when the *csnk-1* RNAi-treated PCMD-1-PBD^m^ PCM dissembled and dissolved extremely rapidly (Figure 4 C). It almost seemed to be torn apart by the exerted pulling forces, showing a significantly lower value of circularity and increased area than the PCMD-1^WT^ control (Figures 4 D and E).

In summary, microtubule pulling forces are not the only cause for the shape distortion of the maturing PCM scaffold in PCMD-1-PBD^m^ at prometaphase and metaphase and their reduction only partially restores the shape of a disassembling PCM in anaphase. *csnk-1(RNAi)* experiments revealed that the distorted PCM scaffold in PCMD-1-PBD^m^ mutant is extremely labile and unable to withstand an increase in microtubule pulling forces.

### Reducing PCMD-1 levels leads to a compromised PCM scaffold formation

We found centrosomal PCMD-1 levels to be significantly decreased in the PCMD-1- PBD^m^ embryos (Figure 1C and D), and this decrease could directly or indirectly cause the distortion of the PCM scaffold. To test this possibility, we decreased PCMD-1 levels by treating worms expressing the GFP-tagged PCMD-1^WT^ construct with RNAi against GFP (Figure S4 A). Upon RNAi treatment mean PCMD-1 levels at the centrosome dramatically decreased to 12.6% of the mock treated controls (Figure S4 B). These levels were lower than the decrease caused by the mutations in PCMD-1-PBD^m^. The PCM mass was comparable to the mock treated embryos (Figure S4 C), however, similar to the PCMD-1-PBD^m^, the PCM scaffold appeared more disorganized (Figures S4 D and E). This indicates that low PCMD-1 levels at the centrosome causes PCM disorganization. To better illustrate how the degree of PCMD-1 levels relates to the PCM scaffold circularity in RNAi-treated PCMD-1^WT^ and PCMD-1-PBD^m^ embryos, we individually plotted all datapoints on a scatterplot (Figure S4 F). For mock-treated or untreated PCMD-1^WT^ embryos mean circularity values were close to 0.9 (gray and black dotted line). In GFP RNAi-treated PCMD-1^WT^ controls mean circularity dropped just below 0.8 (green dotted line). Interestingly, although the centrosomal intensities of PCMD-1-PBD^m^ were less strongly decreased than the ones of GFP RNAi-treated PCMD-1^WT^ embryos, the PCM appeared more distorted, with mean circularity values dropping to 0.7. We would like to point out that some of the PCMD-1-PBD^m^ centrosomes (Figure S4 F, blue dots circled with black line) with medium PCMD-1 levels showed a very high degree of PCM disorganization.

In summary, lowering PCMD-1 levels at the centrosome can destabilize the PCM scaffold. However, the degree of disorganization in PCMD-1-PBD^m^ exceeds the one caused by a simple decrease in PCMD-1 levels, pointing towards an additional layer of PCM scaffold stabilization by PCMD-1.

### PBD-sites in the N-terminal part of PCMD-1 are needed to form a stable PCM scaffold

To further discriminate between these two levels of regulation and to narrow down the region playing a role in PCM stabilization, we set out to create a mutant version of PCMD-1 which would uncouple its function in centrosome destabilization from centrosome localization. In our previous studies we have shown that the C-terminal part of PCMD-1 is needed for its robust loading to the centrosome (Stenzel et al., 2021). We reasoned that mutations in the C-terminal part of the protein could be responsible for low PCMD-1 levels at the centrosome and swapping the mutant C-terminus with a wild type transgene would potentially lead to increased PCMD-1 levels at the centrosome. Therefore, we created a transgene harboring mutated PLK-1 binding sites only in the N-terminal part (PCMD-1-PBD^m^ N^ter^) and a complementary transgene, carrying mutations in the C-terminal part (PCMD-1-PBD^m^ C^ter^) (Figure 5A). Indeed, by reintroducing wild type binding sites in the C-terminal part of the protein, we could significantly elevate centrosomal PCMD-1 levels by restoring levels up to 50% of the control strain in PCMD-1-PBD^m^ N^ter^ (Figures 5 B and C). Interestingly, although protein levels were comparable to PCMD-1^WT^ (Figure S1C), centrosomal levels in the PCMD-1-PBD^m^ C^ter^ transgene were only rescued to about 76% of the control, suggesting a role of the C-terminal PLK-1 binding sites in PCMD-1 localization and PCM recruitment. However, this difference in centrosomal levels did not affect PCM scaffold shape dynamics over time. Mean PCM area and circularity of PCMD-1- PBD^m^ C^ter^ at prometaphase, metaphase and anaphase where not different from control embryos (Figures 5 D, E and S5 A).

**Figure 5:**
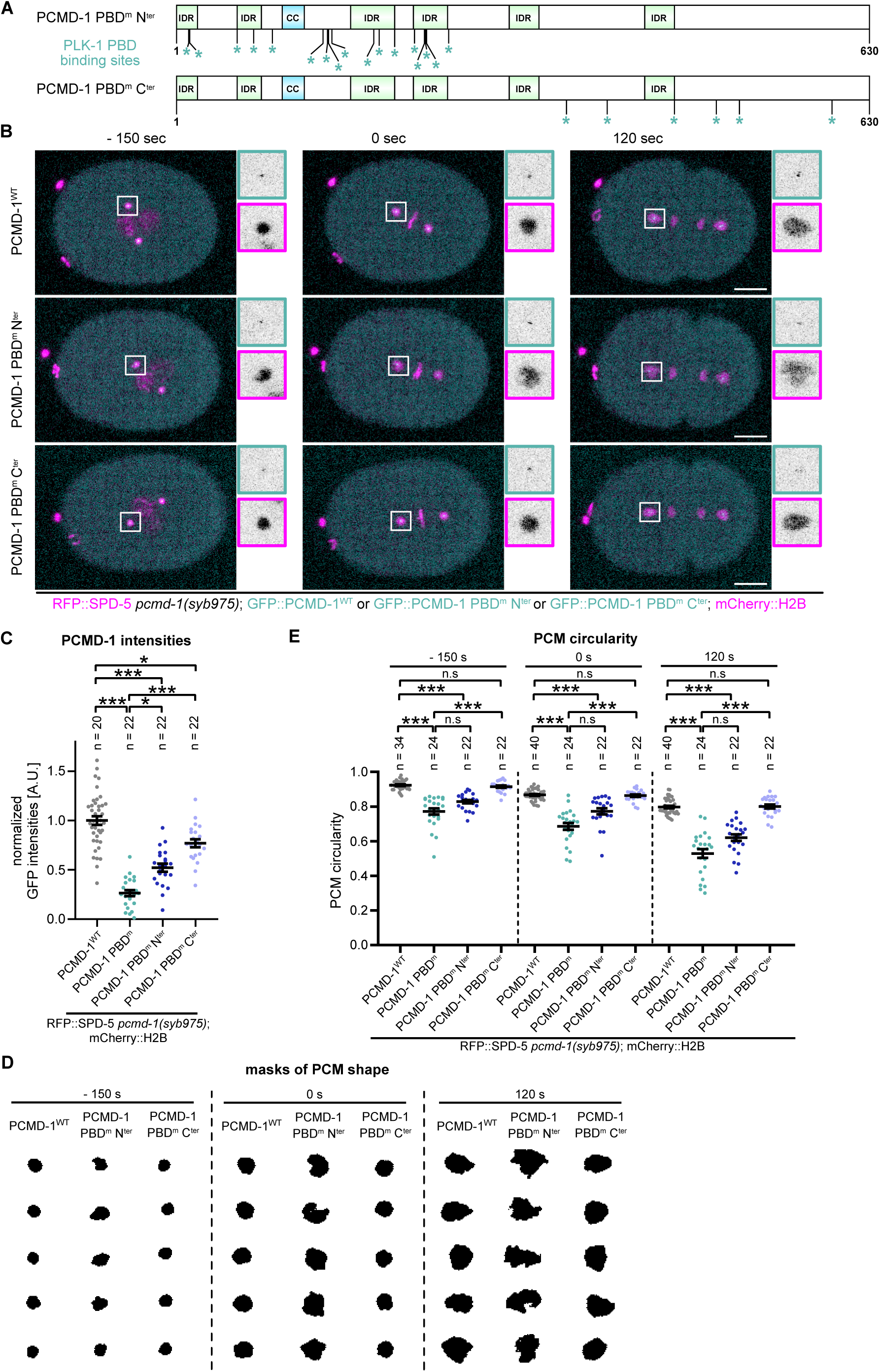
PBD-sites in the N-terminal part of PCMD-1 are needed to form a stable PCM scaffold. A) Schematic representation of the domain structure of PCMD-1 PBD^m^ Nter and PCMD-1 PBD^m^ Cter, with predicted PLK-1 PBD^m^ binding sites indicated by an asterisk under the protein structure. B) Stills of timelapse movies of embryos expressing GFP::PCMD-1^WT^, GFP::PCMD-1 PBD^m^ N^ter^ and PCMD-1 PBD^m^ C^ter^ in combination with mCherry::H2B and RFP::SPD-5 in the *pcmd-1(syb975)* background. Centrosomal areas are shown enlarged in insets. Scale bar = 10 μm. C) Normalized GFP intensities at the centrosome during metaphase. p-value from an Anova with subsequent TukeyHSD. D) Representative examples of PCM shapes of GFP::PCMD-1^WT^, PCMD-1 PBD^m^ N^ter^ and PCMD-1 PBD^m^ C^ter^ at indicated timepoints. E) Quantification of PCM circularities at indicated time points. p-values from an Anova with subsequent TukeyHSD and Kruskal Wallis test with Dunn post-hoc. For all movies: 0 sec is metaphase. n = number of analyzed centrosomes.

In contrast, in PCMD-1-PBD^m^ N^ter^ embryos the PCM scaffold still appeared very labile and deformed (Figures 5 D, E and S5 A). The PCM disorganization started from prometaphase on and increased at metaphase and anaphase. Mean PCM circularity and area at prometaphase, metaphase and anaphase were still comparable to the ones of PCMD-1-PBD^m^.

To mimic centrosomal PCMD-1 levels comparable to PCMD-1-PBD^m^ N^ter^, we subjected the control PCMD-1^WT^ strain to a weak RNAi against GFP (Figure S5 B). Mean PCMD-1^WT^ levels at the centrosome decreased to 40% of mock treated embryos (Figure S5 B and data not shown). To better visualize the impact of intermediate PCMD-1 levels on PCM disorganization, we plotted centrosomal PCMD-1 levels against PCM disorganization for individual datapoint in the weak GFP RNAi-treated and PCMD-1-PBD^m^ N^ter^ embryos (Figure S5 B). For mock-treated or untreated PCMD- 1^WT^ embryos mean circularity values were again close to 0.9 (gray and black dotted line). In weak GFP RNAi-treated PCMD-1^WT^ embryos mean circularity dropped just below 0.9 (green dotted line). Instead for PCMD-1-PBD^m^ N^ter^ embryos mean circularity dropped even further, below the 0.8 mark (blue dotted line).

Taken together, this data supports the previous suggested two layers of PCM dysregulation in PCMD-1-PBD^m^ and PCMD-1-PBD^m^ N^ter^. One that is caused by the decreased levels of PCMD-1 at the centrosome and a second one, which is induced by the mutations in potential PLK-1 binding sites.

### PCM fragmentation caused by SPD-5(Δ734-918) does not affect PCMD-1 recruitment to the centrosome

While PCMD-1 recruits and tethers SPD-5 to the centrioles, in turn SPD-5 can also recruit PCMD-1 to the ciliary base in positive feedback loop (Garbrecht et al., 2021). Therefore, a labile PCM scaffold could potentially impair PCMD-1 recruitment to the centrosome and be an indirect cause of lowered PC PCMD-1-PBD^m^ levels at the centrosome. To test this possibility, we used the Δ734-918 deletion in the C-terminal ‘coiled-coil-long’ domain in the endogenously tagged SPD-5 protein (Rios et al., 2024).

In this mutant background the PCM exhibits microtubule dependent PCM fragmentation, due to an aberrant self-interaction of SPD-5. We combined this strain with an endogenously GFP-tagged PCMD-1 and tested whether PCMD-1 levels were affected in this background.

As shown in Rios et al (2024), in SPD-5(Δ734-918) mutant embryos the PCM mass is largely diminished compared to control embryos, and the PCM scaffold has an irregular and distorted shape (Figures S6 A, C and D). In prometaphase, when PCMD- 1-PBD^m^ is already disordered (compare Figure 1 G to S6 D), SPD-5(Δ734-918) PCM scaffold appeared spherical and comparable to the control. Deviations in mean circularity of SPD-5(Δ734-918) PCM became apparent at metaphase, and increased during anaphase, coincidental with the increased tension exerted by the microtubule pulling forces (Figures S6 C and D). At metaphase mean circularity values dropped from 0.9 to 0.7 in the SPD-5(Δ734-918) embryos. Despite a less dense and more disorganized PCM in SPD-5(Δ734-918) animals, PCMD-1 levels were similar to the ones of a wild-type SPD-5 PCM. Therefore, a disorganized PCM does not lead to failed PCMD-1 recruitment to the centrosome.

Cumulatively these results suggest that appropriate PCMD-1 levels at the centrosome are needed to stabilize the PCM scaffold. However, PCMD-1 levels are not sufficient to explain the disorganization. Additional functional changes due to the introduced mutations at PBD sites in the N-terminal part of PCMD-1 are accountable for PCM disorganization.

### Centrosome separation is delayed in PCMD-1-PBD^m^ N^ter^

In vertebrates, centriole separation depends on the cleavage of pericentrin by separase at anaphase onset (Kim et al., 2015; Lee and Rhee, 2012; Matsuo et al., 2012). In *C. elegans* separase does not play a role in centriole separation. According to the current model pericentriolar material traps the centriole pair at the spindle pole (Cabral et al., 2013). At anaphase PCM weakening and disassembly by microtubule pulling forces allows centrioles to separate. Whether this release is a passive process and merely a consequence of PCM disassembly or alternatively, requires a relay of microtubule pulling forces to the centrioles is not known. The PCM scaffold in PCMD- 1-PBD^m^ N^ter^ is weakened and labile and is therefore an ideal setting to test whether a compromised PCM can still relay the tension generated by the pulling forces to the centrioles.

Therefore, we set out to measure and analyze centriole separation dynamics in a three-dimensional volume at each spindle pole in PCMD-1-PBD^m^ N^ter^ embryos (Figure 6 A) as the intermediate PCMD-1 levels allow us to track the centrioles easy over time. In control embryos, starting from 60 sec after metaphase two slightly separated, but still largely overlapping foci were evident. We consider this time period as ‘centriole disengagement’. On average, the first timepoint at which one could clearly discriminate two independent centriole foci was at 150 sec for the anterior pole and 120 sec for the posterior. We set the distance of 0,6 μm as a threshold and determined this timepoint as the ‘onset of centriole separation’ (Figures 6 B and C). We scored centriole separation until the newly formed centrosomes touched the nuclear envelope, where other dynein dependent mechanism could influence centriole movement (De Simone et al., 2016). Centriole separation dynamics differs at the two poles, due to the differences in microtubule pulling forces and PCM disassembly dynamics. At the anterior centrioles separate slower, with a flatter curve over time. Instead, at the posterior pole, where the microtubule pulling forces are higher, centriole separation started faster and slightly plateaued around 300 sec after metaphase (Figures 6 B and C).

**Figure 6:**
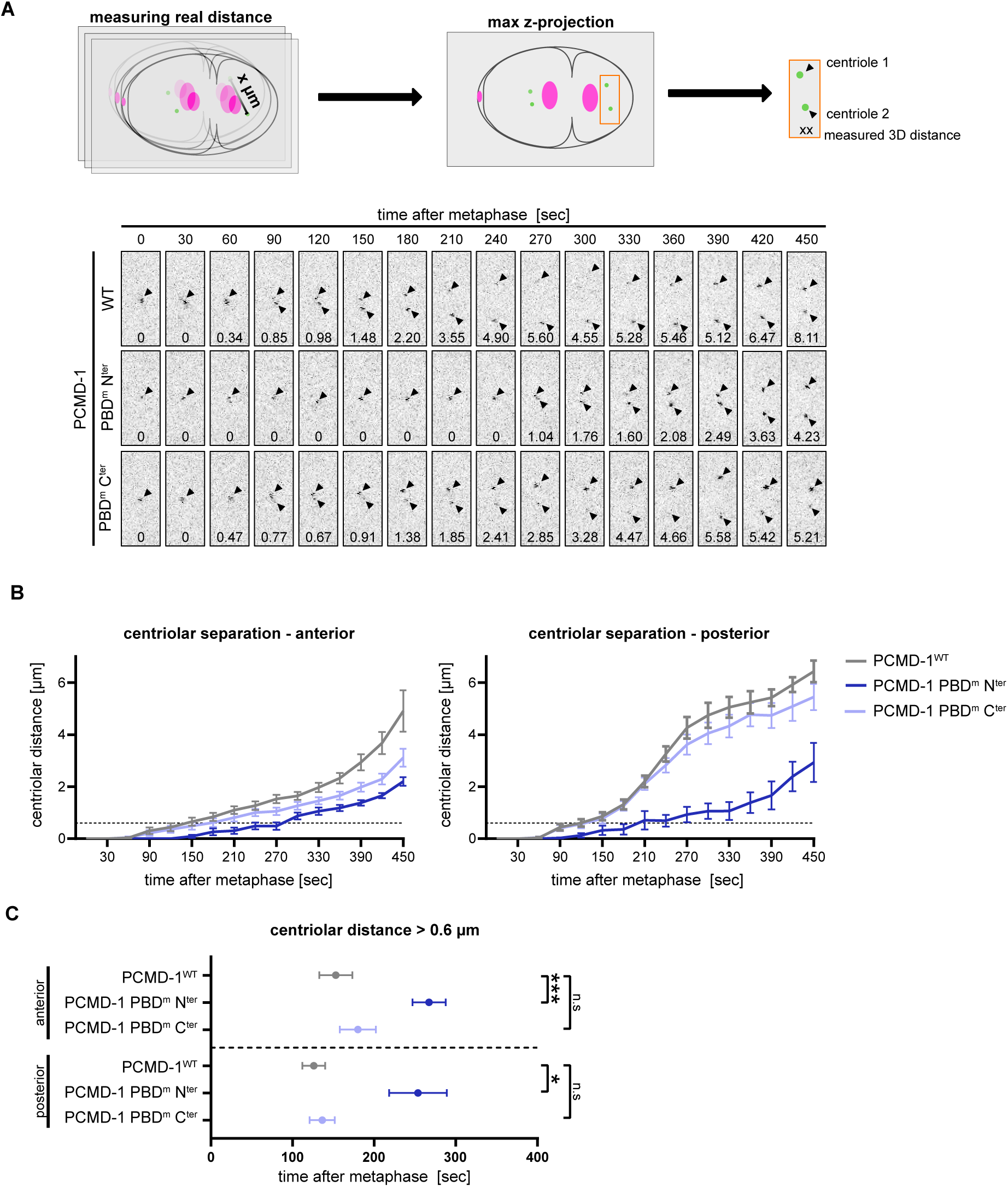
Centrosome separation is delayed in PCMD-1-PBD^m^ N^ter^. A) Schematics of 3D measurement of centriole separation. Analysis was performed at the anterior and posterior spindle pole. Representative timelapse images of centriole separation at the posterior pole of GFP::PCMD-1^WT^, GFP::PCMD-1 PBD^m^ N^ter^ and PCMD-1 PBD^m^ C^ter^. B) Quantification of the centriolar distance during the first cell division at the anterior and posterior spindle pole of GFP::PCMD-1^WT^ (n=10, in grey), GFP::PCMD-1 PBD^m^ N^ter^ (n=11, in violet) and GFP::PCMD-1 PBD^m^ C^ter^ (n=11, in blue) embryos. For all movies: 0 sec is metaphase. Bold lines represent the mean +/- SEM. Dashed lines indicates centrioles at a distance of 0.6 µm, a threshold defined as the onset of visible centriole separation. C) Quantification of the time point of onset of centriolar separation at the anterior and posterior spindle poles, defined as a distance of 0.6 µm. p-values from Kruskal-Wallis test with Dunn post-hocand an Anovawith subsequent TukeyHSD. n = number of analyzed centrosomes.

In PCMD-1-PBD^m^ N^ter^ embryos centriole separation was significantly delayed. Two distinct foci could be first discriminated at 270 sec at the anterior and at 240 sec at the posterior pole (Figures 6 A (lower panel), B and C). The two foci remained in close proximity for a longer period of time and eventually moved apart, but within the timeframe of our analysis, they never reached the same distance as the control embryos. Interestingly, the posterior pair separated with a similar dynamic as the pair at the anterior pole. In contrast, in PCMD-1-PBD^m^ C^ter^, where the PCM is not disorganized, centrioles separated similar to the control ones (Figures 6 A (lower panel), B and C). Centrioles disengagement and separation started around the same time as in control embryos. The general separation dynamics were similar to the control, with the exception that at the anterior pole, separation was somewhat delayed at later timepoints.

This data speaks for a model where a strong and ductile PCM scaffold relays the microtubule pulling forces to the centrioles during separation. If so, we would expect that any kind of weakening of the PCM should have a similar effect on centriole separation.

To test this idea, we once again turned to the SPD-5(Δ734-918) mutant embryos and analyzed centriole separation in this background (Figures S6 E). To our surprise we found that centriole separation was largely unaffected. Mean timepoints of the onset of centriole separation at both poles coincided with the one of the control embryos and separation dynamics was comparable to controls. Only at the posterior pole centrioles separated with a higher variability and on average did not separate to the same extent as in the control embryos.

In summary, at the poles of PCMD-1-PBD^m^ N^ter^ embryos centrioles fail to separate in a timely manner and the delay in separation cannot be explained solely by the disorganization of the PCM scaffold. Mutated PBD-binding sites in PCMD-1 could act as a part of the relay mechanism by which centrioles are separated during PCM disassembly.

## DISCUSSION

Here we show, that PCMD-1 modulates PLK-1 levels at the centrosome and contributes to the PCM integrity and centriole separation.

Over the course of mitosis, the PCM scaffold isotopically grows and undergoes changes in its physical properties (Laos et al., 2015; Mittasch et al., 2020). It transitions from a strong and ductile state, resisting microtubule pulling forces at metaphase, to a labile or more brittle state in anaphase, where it undergoes dispersal and dissolution (Mittasch et al., 2020). Interestingly, recent cryo-ET analysis could not reveal any structural differences of the PCM over the cell cycle (Tollervey et al., 2024), suggesting that the described alterations in physical properties are driven by qualitative modulations of PCM components. The switch to a brittle scaffold that is less resistant to microtubule forces and prone to disassembly, is taking place concomitant with the drop of PLK-1 centrosomal levels, and chemical inhibition of PLK-1 at metaphase triggers premature PCM disassembly (Cabral et al., 2019; Mittasch et al., 2020). It has been proposed that the balance between PLK-1 phosphorylation and counteracting dephosphorylation by the counteracting PP2A phosphatase drives the abovementioned changes in material properties of the PCM (Magescas et al., 2019; Mittasch et al., 2020).

In our previous work we showed that PCMD-1, a protein with several intrinsically disordered regions (IDRs), interacts with SPD-5 and PLK-1 and that PCMD-1 is able to recruit these two proteins to an ectopic cellular location (Stenzel et al., 2021). However, whether this interaction is needed to maintain PLK-1 at the centrosome and how failure to do so affects PCM scaffold dynamics, was not explored. By mutating putative PLK-1 binding sites in PCMD-1, we created a situation in which wild type centrosomal PLK-1 levels are not reached at metaphase. Due to the close vicinity of the centrosome to the nuclear membrane, we were unable to reliably measure exclusively centrosome levels of PLK-1 at prometaphase, but by projecting the metaphase levels back over time, we speculate that they could be lower as well. This reveals a new function of PCMD-1 in PLK-1 recruitment and maintenance at the centrosome. We found that PLK-1 in turn phosphorylates PCMD-1 *in vitro*, however this phosphorylation seems to play no significant role in PCMD-1 recruitment or its function in forming a functional scaffold. One caveat of our analysis is that we might have missed PLK-1 phosphorylation sites that do not match the known consensus motifs. Intriguingly, by mutating 30 potential PLK-1 phosphorylation sites, which makes up nearly 5% of the entire protein, we did not affect its overall dynamics and function. This most likely is explained by the flexible nature of PCMD-1, which has no predicted functional domains.

In contrast to PLK-1, SPD-5 is recruited to the centrosome to an almost wild type level in PCMD-1-PBD^m^. Nevertheless, the scaffold still assembles in an aberrant manner. A distorted and flared PCM scaffold is already evident during PCM maturation when the cell transitions from prophase to metaphase. Can the PCM disorganization be attributed the slightly decreased PCM mass recruitment?

Compared to PCMD-1-PBD^m^, in SPD-5(Δ734-918) the PCM mass is much lower, indicating a less efficient scaffold recruitment. Nevertheless, the low amount of PCM mass does not translate into a higher degree of disorganization at prometaphase, arguing that the relatively small change in PCM mass in PCMD-1-PBD^m^ compared to PCMD-1^WT^, is not the main driver of failed PCM scaffold assembly. Interestingly, in contrast to PCMD-1-PBD^m^ the shape of the SPD-5(Δ734-918) scaffold is not altered during prometaphase. It seems like PCM maturation of the SPD-5(Δ734-918) scaffold is unaffected, and defects in the PCM scaffold are only reveled at the metaphase when microtubule pulling forces become most active and displace the spindle. In contrast, in PCMD-1-PBD^m^ the scaffold seems to assemble in an abnormal way from the moment of its recruitment to the centrioles.

How aberrant the PCM scaffold actually is, is best described by its response to the modulation of microtubule puling forces. PCM distortion at prometaphase and metaphase is evident by exaggerated deformation of its shape parameters, when higher pulling forces are applied under *csnk-1 RNAi.* In contrast, when pulling forces are decreased by using *gpr-1/2 RNAi,* the shape of the distorted centrosome is not restored to wild type circularity. Although we cannot entirely rule out that residual microtubule pulling forces act on the PCM, PCM scaffold shapes do not change significantly, despite the failed posterior displacement of the metaphase spindle, which is a readout for the decreased of microtubule pulling forces at this timepoint. Together this indicates that the PCM in PCMD-1-PBD^m^ is formed in a less condensed manner and the initial shape distortion of the PCM cannot entirely be attributed to microtubule pulling forces, and is most likely a result of an intrinsically aberrant assembly of the scaffold. In anaphase the already structurally weakened PCM is completely unable to counteract increased microtubule pulling forces. The scaffold is literally torn apart and dispersed. The disorganized PCM scaffold in prometaphase very much resembles the one caused by centriole ablation before NEBD (Cabral et al., 2019). Due to the ablation, the PCM scaffold seems to have lost its tether at the center and disperses in a microtubule dependent manner. However, in contrast to centriole ablation, in PCMD- 1-PBD^m^, reduction of microtubule pulling forces does not restore PCM circularity, supporting the possibility that the PCM scaffold has intrinsically different properties from the moment of its formation.

By modulating wild type PCMD-1 levels by RNAi and by mutating the PBD-binding sites only in the N-terminus of PCMD-1, we could reveal a two-tier regulation of scaffold shape and integrity. The first tier is explained by the reduced level of PCMD- 1 at the centrosome. However, by minimizing PCMD-1 at the centrosome, only an intermediate degree of PCM disorganization can be achieved. The extremely high degree of PCM distortion in PCMD-1-PBD^m^ and PCMD-1-PBD^m^ N^ter^ can only be explained by the additional impact of the mutated binding sites.

The changes in PCM properties allows us for the first time to test the model put forward for centriole separation in *C. elegans* (Cabral et al., 2013; Magescas et al., 2019). According to the current model centrioles are trapped in the PCM and upon PCM disassembly in late anaphase centrioles are passively released and separated. According to this model we would expect centrioles embedded in a softer and more labile PCM to separate prematurely. Instead, our measurements in PCMD-1-PBD^m^ showed a delay in centriole separation at both poles. This can be explained by two mutually nonexclusive mechanisms. First, PCMD-1, similar to the human PCNT, could act as a linker and this linker would require PLK-1 binding for a timely cleavage. Failure in proper PLK-1 recruitment and binding at metaphase could delay the process of separation in late anaphase. Alternatively, after PCM disassembly in late anaphase, timely centriole separation could require a direct tether or relay mechanism that transmits the forces generated by the astral microtubules to the centrioles in the moment of PCM rupture. In PCMD-1-PBD^m^ this relay mechanism could be dysfunctional, allowing for PCM to be torn apart easier, while leaving behind the centrioles to be separated by their own slow kinetics (Figure 7).

**Figure 7:**
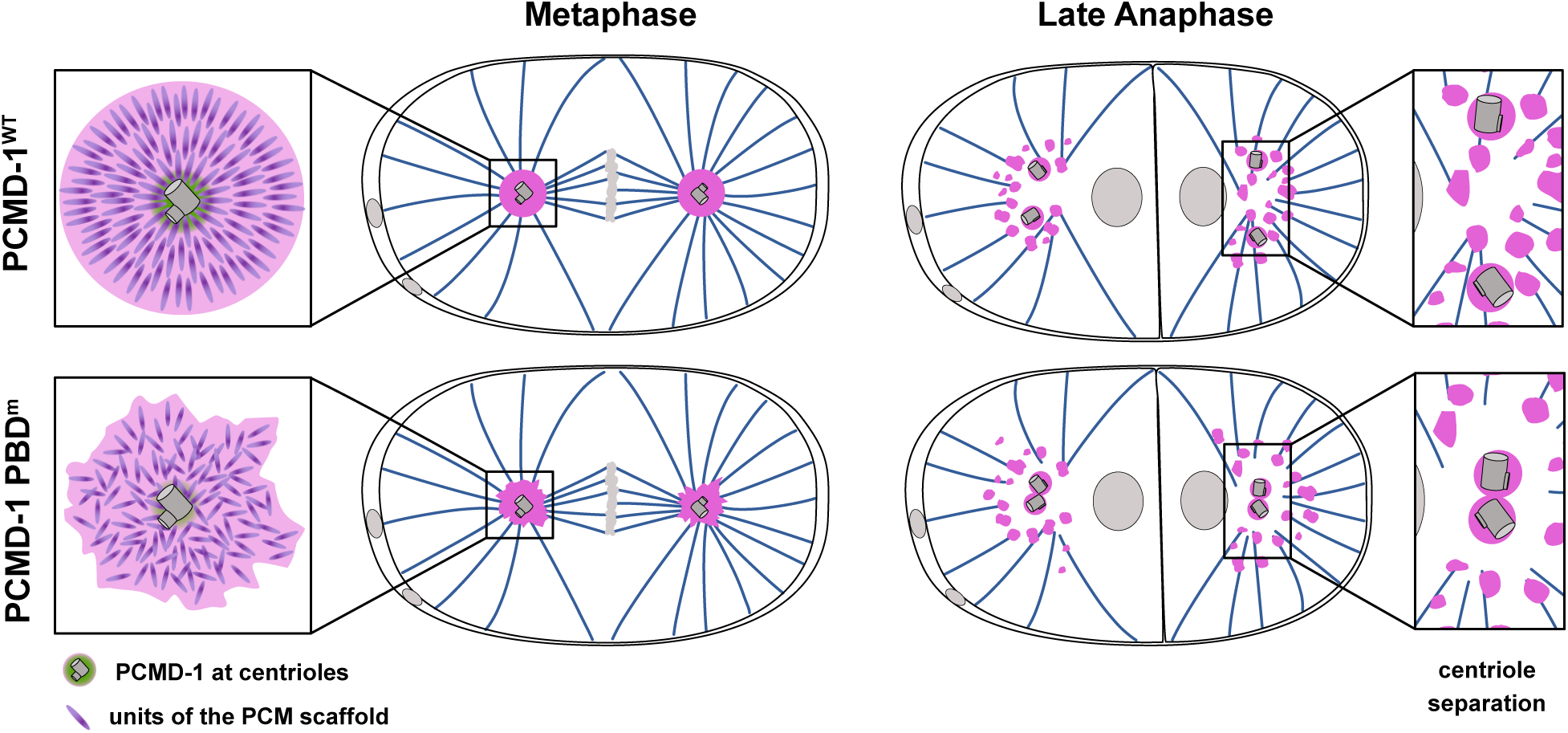
Model of PCM scaffold destabilization and delayed centriole separation in PCMD-1 PBD^m^ embryos. Left panels: Close to the centrioles, PCMD-1^WT^ together with PLK-1 (not shown) biases the multimodal self-interactions of SPD-5 towards a certain intrinsic order. A rotational symmetry propagates this order along the entire PCM scaffold, giving the PCM strength and integrity. PCMD-1 PBD^m^ fails to bias the macromolecular interactions during PCM seeding and maturation, leading to a scaffold formation by randomized macromolecular self-interaction, which in turn leads to a destabilized PCM. Right panels: After PCM disassembly in late anaphase, pulling forces from astral microtubule are relayed to the centrioles, allowing for a timely centriole separation. In PCMD-1-PBD^m^ the disorganized and labile PCM is torn apart easier, leaving behind the centrioles to separate by their own slow kinetics in absence of a functional relay mechanism. Note: centrioles and the PCM scaffold units are not drawn to scale.

We cannot exclude that by mutating putative PLK-1 binding sites in the PCMD-1 we alter its interaction with SPD-5. PCMD-1 is an intrinsically disordered protein, the structure of which is poorly predicted and can change upon binding with different partners or in response to posttranslational modifications. In fact, when expressed in HEK293T cells, PCMD-1-PBD^m^ is unable to recruit SPD-5, pointing towards an altered interaction of the two proteins. In contrast, in *C. elegans* embryos SPD-5 is recruited to the centrosome however an altered binding mode could still undermine scaffold stability independent of its recruitment or levels.

How could PCMD-1, which is mostly concentrated at the centrioles, affect integrity of a micron scale structure? We previously used super resolution microscopy to describe low or close-to-detection-limit signals of PCMD-1 along SPD-5 fibers, which could potentially stabilize the entire scaffold (Erpf et al., 2019). Alternatively, we propose that PCMD-1 together with PLK-1 and SPD-5 forms a complex close to the centrioles, that facilitates PCM maturation and biases the PCM to assemble into a strong and ductile scaffold. The centrosome was speculated to behave as a non-Newtonian liquid, such as a liquid crystal with physical properties that fall between ordinary liquids and solids (Woodruff, 2021). Properties of a liquid crystal are depended on the orientation of its units or molecules. If so, one could envision that PCMD-1 seeds the PCM scaffold in close vicinity of the centrioles by bringing together SPD-5 and PLK-1. At the centrioles, PCMD-1 and PLK-1 would bias the multimodal self-interactions of SPD-5 towards a certain intrinsic order (Rios et al., 2024). A rotational or translational symmetry could propagate this order along the entire PCM scaffold, giving the PCM strength and integrity (Figure 7). The failure to bias the macromolecular interactions during centrosome seeding and maturation, be it due to lowered PLK-1 levels, aberrant SPD- 5 binding or a combination of both, could result in a scaffold built by randomized macromolecular self-interaction, which in turn leads to a destabilized PCM with more liquid-like properties.

In summary, our work demonstrates that PCMD-1 plays a stabilizing role during PCM scaffold formation and that its function is needed for timely centriole separation at late anaphase.

## ACKNOWLEDGMENTS

We thank the CALM facility for support during live-cell imaging. We like to express our deep gratitude to Monica Gotta for sharing the unpublished strain expressing *plk- 1(syb2781)::mCherry,* to Lionel Pintard for sharing the cePLK-1 Kinase, Saskia Hütten for the support with protein purification, and to Jeff Woodruff for sharing the strain expressing *spd-5(wow36_syb5737[D734-918])*. Some strains were provided by the CGC, which is funded by NIH Office of Research Infrastructure Programs (P40 OD010440). We would also like to thank master students Felicia Haxel, Disha Shenai, Sasha Degtyareva and Alex Schilder for support with building the worm strains and conducting the experiments.

## AUTHORS CONTRIBUTIONS

A.S. conducted most experiments and analyzed most data of the manuscript; she was helped by A.H. and F.W.; all statistical analysis were run by A.S.; A.S. and T.M-D., in discussions with E.Z. conceived the project and wrote the manuscript.

## MATERIALS AND METHODS

*C. elegans* strain maintenance and generation
*C. elegans* strains (Table 2) were maintained under standard conditions on NGM plates seeded with *E. coli* (OP50) at 15°C(Brenner, 1974). If not indicated otherwise, worms were shifted to 25°C at the late L3-L4 stage for 16-20 h for experimental usage. Transgenic worm strains were generated by single-copy integration of the respective transgenes using the MosSCI system (Frokjaer-Jensen et al., 2008). Transgenes were cloned into the pCFJ350 vector and injected into the EG6699 (chromosome II).

### Prediction of PLK-1 phosphorylation or PBD-binding sites of PCMD-1

PLK-1-PBD binding sites of the of PCMD-1 were predicted by searching for the consensus Polo-binding motif S-S/T as described in (Alvarez-Rodrigo et al., 2019) and by using the GPS-PBS (Guo et al., 2020) and the GPS-Polo (Liu et al., 2013) prediction programs with a medium threshold. All predicted sites were selected and mutated.

Potential PLK-1 phosphorylation sites were identified using the kinase-specific phosphorylation prediction system GPS 5.0 with a medium threshold (Wang et al., 2020). All predicted sites were mutated.

### Embryonic survival assay

L4 worms were singled and incubated at 25 °C for 16-20 hours. The laid eggs were counted and compared to the hatched larvae after 48 hours to assess the embryonic lethality. Control groups for Figures S1B and S2D are the same.

### RNA-mediated interference

RNAi experiments against GFP and *gpr-1/2* were performed after feeding worms with the bacterial clones L4440_GFPi and III-5C03, respectively for 16-20 hours at 25°C(Kamath et al., 2003). Weak GFP RNAi condition was achieved by mixing GFP RNAi bacteria with bacteria carrying the empty pL4440 vector 1:4. For RNAi against *csnk-1*, a L4440 vector was generated carrying the CDS of *csnk-1,* transformed into *E. coli* HT115(DE3) bacteria and worms were fed for 48 hours at 25 °C. Bacteria carrying the empty pL4440 vector were used as negative control (mock).

### Microscopy

Young adult worms were immobilized and dissected in 4µl 2.5 µM Levamisol in M9 buffer to extract the embryos and mounted on 4 % agarose pads. Live-cell imaging was performed on an inverted Leica Stellaris 5 Point Scanning Confocal Microscope with a resolution of 512x512 pixels using an 100X HC PL APO oil immersion objective and controlled by LAS X software. *z*-stacks were taken every 30 s with a step size of 0.7 µm and 2×2 binning. For representative images from one-cell embryos max. z-projections of a fixed volume of 20 slices were generated.

### Protein overexpression and purification

MBP-PCMD-1-6xHis and MBP-6xHis proteins were expressed in BL21(DE3) bacteria. At an OD600 of 0.4 – 0.6 the bacterial culture was subjected to cold-shock and protein expression was induced for 20 hours at 17 °C with 0.1 mM IPTG. Bacterial pellets were harvested, washed with cold PBS and snap frozen in liquid nitrogen. 1g of bacterial pellet was resuspended in 5 ml lysis buffer (20 mM Tris-HCl pH 7.5, 500 mM NaCl, 1 mM DTT, 0.2 % Triton X-100; supplemented with Complete Mini Protease inhibitor (Roche, 11836153001)) and lysed by sonification. Cleared lysates were incubated with Ni-NTA agarose (QIAGEN, 30210) for 1 hour. Agarose beads were washed three times (50 mM Tris-HCl pH 7.5, 500 mM NaCl, 50 mM Imidazol, 10 % Glycerol) with 10 column volumes, the second wash included high NaCl concentration of 1 M. Protein was eluted (50 mM Tris-HCl pH 7.5, 500 mM NaCl, 500 mM Imidazol, 10 % Glycerol) and buffer exchange (25 mM Tris-HCl pH 7.5, 250 mM NaCl, 25 mM Imidazol, 5 % Glycerol) was performed for three times while protein concentration (Merck, UCF8100) took place. Concentrated protein was eluted and stored at -80 °C until further usage.

### Kinase assay

Kinase assays were performed as in (Schneid et al., 2021) using ∼300 ng MBP- PCMD-1-6xHis or MBP-6xHis and ∼200 ng *C. elegans* PLK-1 kinase (kind gift from Lionel Pintard) in kinase buffer (50 mM Tris-HCl pH 7.5, 10 mM MgCl_2_, 135 mM Saccharose, 0.01 % Triton, 1 mM DTT).

### Immunoblotting

For immunoblotting a fixed number of worms was picked in M9 buffer and washed four times. Buffer volume was reduced and respective amount of 2x Laemmli buffer was added. Samples were incubated at 95 °C for 10 minutes, sonicated for 10 minutes and again incubated at 95 °C for 10 minutes. After applying samples on an SDS-page and Western Blotting, PCMD-1 transgene proteins with GFP-tag were probed with a GFP antibody (mouse monoclonal 1:600, Roche) or α-Tubulin antibody (mouse monoclonal 1:7500, Sigma-Aldrich). Proteins of interest were detected using HRP-conjugated secondary antibodies against mouse (1:7500, Bio-Rad Laboratories). The proteins were visualized using Amersham ECL Prime Western Blotting Detection Reagent (GE Healthcare).

### Cell cycle length measurements

Young adult worms were immobilized and dissected in 4µl 2.5 µM Levamisol in M9 buffer to extract the embryos and mounted on 4 % agarose pads. Live-cell imaging was performed on a Zeiss Axio Imager.M2 and ‘Time to Live’ software. Every 35 s Z-stacks were acquired through the embryos using 0.7 μm step size in a total volume of 25 µm, from pronuclear migration until end of second cell cycle. Time from NEBD in the one-cell embryo until NEBD in the AB cell was measured.

### Measuring spindle positioning

Spindle position was measured in Fiji (Schindelin et al., 2012) at metaphase by drawing a linescan from the anterior to posterior cortex of the embryo. mCherry-H2B intensities along the linescan were measured to identify the position of the DNA metaphase plate relative to embryo length.

### Fluorescence intensity analysis

GFP, RPF and mCherry intensities were measured on raw images by analyzing *z*- stacks with ManualTrackMate in Fiji (Schindelin et al., 2012; Tinevez et al., 2017). The time point of measurement was determined by the DNA condensation pattern visualized by the mCherry::H2B marker. Fluorescent signals were measured in a sphere with a fixed radius (GFP: 0.782 µm, RFP::SPD-5: 3.328 µm, PLK-1::mCherry: 1.013 µm). A sphere with the same radius was used to measure the cytoplasmic background signal and the background signal outside the embryo. For the each datapoint, the cytoplasmic background signal was subtracted from the fluorescent signal. To take in account the cytoplasmic gradient of PLK-1::mCherry in the one-cell embryo, cytoplasmic background levels were measured in the anterior and posterior side of the embryo and subtracted from the respective centrosomal signal.

### PCM shape analysis

PCM circularity and area were measured at indicated time points. Maximum *z*- projections of the RFP::SPD-5 channel were created and PCM was extracted by removing outliers (radius 2; threshold: 5) and despeckling the whole stack. PCM shape was then converted into black/white outlines using the ‘Otsu white’ threshold. Circularity and area were measured by the Fiji software (Schindelin et al., 2012). Circularity of a shape is defined as 4πx(area/perimeter^2^), whereas 1.0 indicates a perfect circle.

### Measuring centriolar distance

Centriolar distance was measured in 4D stacks after metaphase using the TrackMate Plugin (Ershov et al., 2022) in Fiji (Schindelin et al., 2012). Centrioles were detected using DoG detector with an estimated object diameter of 1 µm, a quality threshold of 5 and subpixel localization. Signals were contrast (0.3 and above) filtered and missed centrioles, especially due to weak signal, were manually added. To track their separation LAP Tracker was used with a Frame-to-frame linking of 10 µm, Track segment gap closing with a distance of 10 µm with maximal 2 frames gap and Track segment splitting with a maximal distance of 10 µm. Tracks were manually adjusted if required, centriolar positions were measured by Fiji and centriolar distance at all analyzed timepoints was calculated by Pythagoras theorem in 3D.

### Cell culture, transfection and imaging

Human embryonic kidney (HEK) 293T cells were cultured in Dulbeccòs Modified Eagle Medium (DMEM) supplemented with 10% fetal calf serum, penicillin (100U/ml) and streptomycin(100µg/ml) at 37°C, 5% CO2. For microscopy the cells were grown to 50-70% confluence on glass coverslips and transfected with expression plasmids using Lipofectamin 2000 (Invitrogen) according to the manufacturer’s instructions. 24h after transfection cells were fixed with 4% paraformaldehyde (15min at room temperature), permeabilized with 1% Triton X-100 in PBS and stained with DAPI (4′,6- Diamidin-2-phenylindol, 10µg/ml). Cells were imaged at the Leica SP5 point scanning confocal microscope with the HCX PL APO Lambda Blue 63x 1.4 oil objective and 512x512 pixel resolution.

### Statistical analysis

The statistical analyses were performed using the software R (Posit team, 2024). RStudio: Integrated Development Environment for R. Posit Software, PBC, Boston, MA. URL http://www.rstudio.com/) and the packages car (Fox and Weisberg, 2019) and FSA (Ogle et al., 2017). Data were tested for normal distribution using the Shapiro-Wilk test and for equal variances using the Levenes Test. Datasets with two groups and parametric conditions were analyzed by Student’s t test, non-parametric conditions by Wilcoxon Rank sum test. For data with multiple groups and parametric conditions an ANOVA was performed and p-values were calculated subsequently with Tukey Honest Significant Differences. For multiple groups and non-parametric conditions, a Kruskal Wallis Rank sum test with Dunn post-hoc was performed. The statistical test used is indicated in the figure legends. In the analysis of 1G/H, 3D/E, 4D/E, 5E, S2G/H, S4D/E, S5A, S6D an independent statistical test was performed for each timepoint. p-values: n.s. > 0.05 > * > 0.01 > ** > 0.001 > ***

## TABLES

Table 1 Mutated sites in PBD and PD

Table 2 *C. elegans* strains used

## VIDEOS

Video 1 PCM scaffold dynamics in PCMD-1^WT^

Video 2 PCM scaffold dynamics in PCMD-1-PBD^m^

Video 3 PCM scaffold dynamics in PCMD-1-PD^m^

## SUPPLEMENTAL FIGURE LEGENDS

**Figure S1: PCMD-1-PBD^m^ leads to failure in PLK-1 and PCM recruitment to microtubules in HEK293T cells**

A) Representative images of EGFP::PCMD-1^WT^, EGFP::PCMD-1 PBD^m^ or cytoplasmic EGFP expressed in combination with mCherry::PLK-1 or mCherry::SPD- 5 in HEK293T cells. Scale bar = 10 μm. n = number of cells; % indicates number of cells with GFP or mCherry at the microtubules. B) Expression levels of indicated constructs in whole worm lysates. Two independent GFP::PCMD-1^WT^ samples are shown. C) Survival rates of GFP::PCMD-1^WT^ and GFP::PCMD-1 PBD^m^ embryos at 25°C. n = number of embryos, p-value from a Kruskal Wallis test with Dunn post-hoc. D) Cell cycle length measured from nuclear envelope break down (NEBD) in the first cell cycle (P0) to the NEBD in the AB cell. p-value from Anova with subsequent TukeyHSD test.

n = number of analyzed embryos.

**Figure S2: Phosphorylation of PCMD-1 by PLK-1 is not required for PCM scaffold integrity**

A) Schematic representation of the domain structure of PCMD-1^WT^, with predicted intrinsically disordered regions (IDR) in green and the coiled-coil region (CC) in blue. Position of the mutated PLK-1 phosphorylation sites is indicated by an asterisk under the protein structure. B) Stills of timelapse movies of embryos expressing of GFP::PCMD-1^WT^ and GFP::PCMD-1 PD^m^ in combination with mCherry::H2B and RFP::SPD-5 in the *pcmd-1(syb975)* background. Centrosomal areas are shown enlarged in insets. Scale bar = 10 μm. C) Coomassie gel and autoradiography of a PLK1 kinase assay with the indicated MBP/6xHis-tagged PCMD-1 fragments. PLK-1* indicates self-phosphorylation of PLK-1. D) Survival rates of GFP::PCMD-1^WT^ and GFP::PCMD-1 PD^m^ embryos at 25°C. n = number of embryos, p-value from a Kruskal Wallis test with Dunn post-hoc. Same control embryos as used in Figure 1C. E) Normalized GFP intensities at the centrosome during metaphase. p-value from a Two Sample t-test. F) Normalized PCM mass at the centrosome during metaphase. p-value from Two Sample t-test. G) Quantification of PCM circularities at indicated time points. p-values from Wilcoxon rank sum test and Two Sample t-test. H) Quantification of PCM area at indicated time points. p-values from Wilcoxon rank sum test and Two Sample t-test.

For all movies: 0 sec is metaphase. n = number of analyzed centrosomes.

**Figure S3: Spindle is positioned centrally when microtubule pulling forces are decreased**

A) Spindle position at metaphase of embryos expressing GFP::PCMD-1^WT^ or GFP::PCMD-1 PBD^m^ in combination with mCherry::H2B and RFP::SPD-5 in the *pcmd- 1(syb975)* background and treated with mock or *gpr-1/2* RNAi. p-values from Anova with subsequent TukeyHSD test. n = number of analyzed embryos.

**Figure S4: Reducing PCMD-1^WT^ levels leads to a compromised PCM scaffold formation**

A) Stills of timelapse movies of embryos expressing GFP::PCMD-1^WT^ in combination with mCherry::H2B and RFP::SPD-5 in the *pcmd-1(syb975)* background treated with mock and GFP RNAi. Centrosomal areas are shown enlarged in insets. Scale bar = 10 μm. B) Normalized GFP intensities at the centrosome during metaphase. p-value from a Welch Two Sample t-test. C) Normalized PCM mass at the centrosome during metaphase. p-value from Wilcoxon rank sum test. D) Quantification of PCM circularities at indicated time points. p-values from Wilcoxon rank sum test and Welch Two Sample t-test. E) Quantification of PCM area at indicated time points. p-values from Two Sample t-test and Wilcoxon rank sum test. F) Scatter plot correlating normalized GFP intensities and PCM circularities at metaphase of PCMD-1 PBD^m^ and GFP::PCMD-1^WT^ treated with mock or GFP RNAi. Dashed line indicates the mean GFP intensity of the corresponding condition. Circled dots highlight datapoints with intermediate GFP::PCMD-1 PBD^m^ levels and high level of disorganization.

For all movies: 0 sec is metaphase. n = number of analyzed centrosomes.

**Figure S5: Correlation of PCMD-1 intensities and PCM scaffold circularities of centrosomes with PBD^m^ N^ter^ and intermediate PCMD-1^WT^ levels**

A) Quantification of PCM areas of GFP::PCMD-1^WT^, PCMD-1 PBD^m^, PCMD-1 PBD^m^ N^ter^ and PCMD-1 PBD^m^ C^ter^ embryos at indicated time points. p-values from Kruskal Wallis test with Dunn post-hoc. B) Scatter plot correlating normalized GFP intensities and PCM circularities at metaphase of PCMD-1 PBD^m^ N^ter^ and GFP::PCMD-1^WT^ treated with mock or weak GFP RNAi. Dashed line indicates the mean GFP intensity of the corresponding condition.

n = number of analyzed centrosomes.

**Figure S6: PCM fragmentation caused by SPD-5(Δ734-918) does not affect PCMD-1 recruitment to the centrosome or centriole separation**

A) Stills of timelapse movies of embryos expressing RFP::SPD-5 or RFP::SPD-5^mut^ (Δ734-918) of embryos expressing endogenously tagged GFP::PCMD-1 in combination with mCherry::H2B. Centrosomal areas are shown enlarged in insets.

Scale bar = 10 μm. B) Normalized GFP intensities at the centrosome during metaphase. p-value from a Two Sample t-test. C) Normalized PCM mass at the centrosome during metaphase. p-value from Two Sample t-test. D) Quantification of PCM circularities at indicated time points. p-values from Two Sample t-test and Welch Two Sample t-test. E) Quantification of the centriolar distance during the first cell division at the anterior and posterior spindle pole of control RFP::SPD-5 (n=10, in grey) and RFP::SPD-5^mut^ (Δ734-918) (n=11, in pink) embryos. Bold lines represent the mean +/- SEM. Dashed lines indicates centrioles at a distance of 0.6 µm, defined as onset of centriole separation. For all movies: 0 sec is metaphase. n = number of analyzed centrosomes.

