## Supplemental Figures and Tables for "PCMD-1 stabilizes the PCM scaffold and facilitates centriole separation"

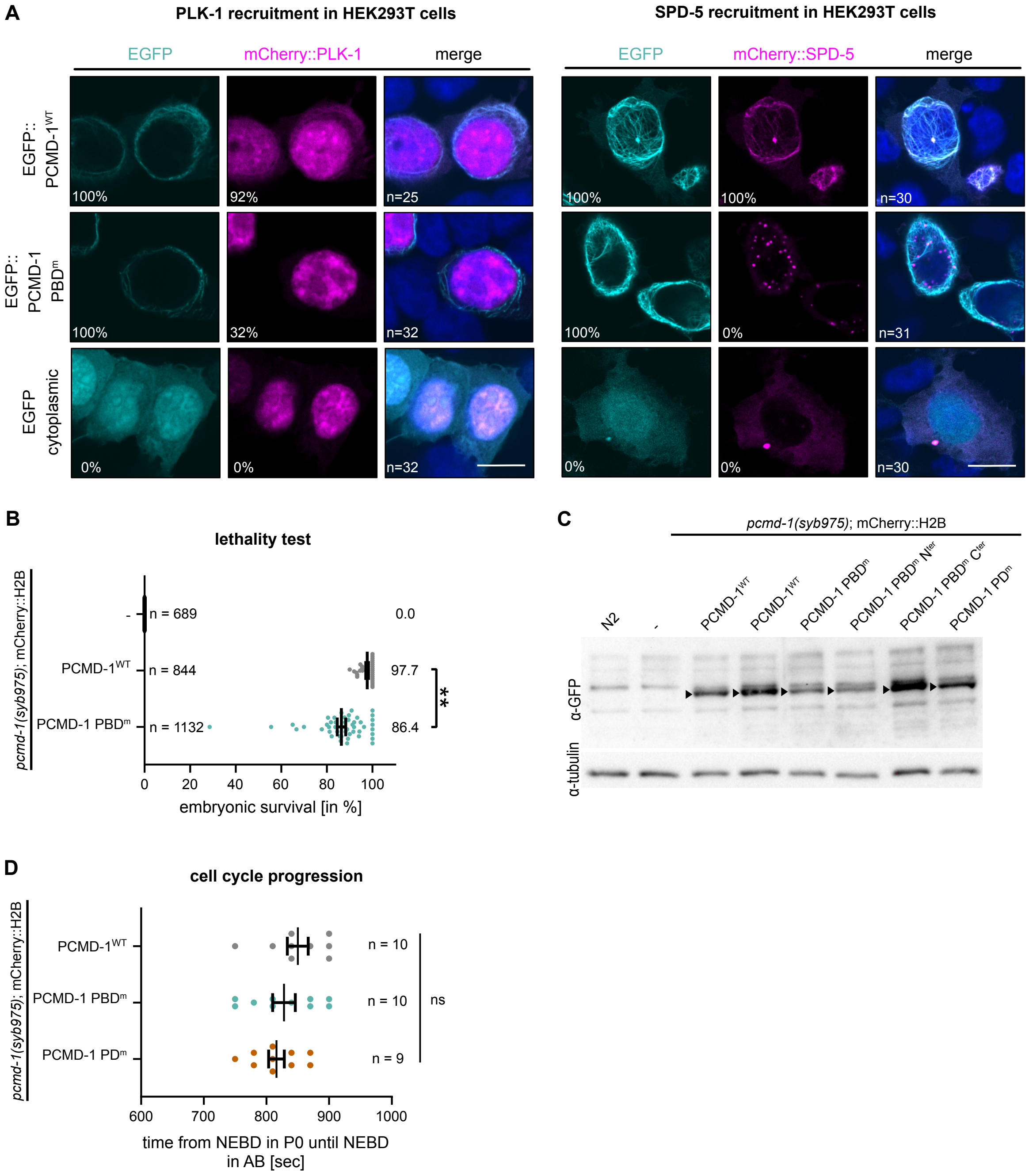
A
PLK-1 recruitment in HEK293T cells
SPD-5 recruitment in HEK293T cells
EGFP
mCherry::PLK-1
merge
mCherry::SPD-5
EGFP:: PCMD-1WT
EGFP:: PCMD-1 PBDm
EGFP cytoplasmic
100%
92%
n=25
32%
n=32
0%
n=30
n=31
B
lethality test
pcmd-1(syb975); mCherry::H2B
PCMD-1WT
PCMD-1 PBDm
n = 689
n = 844
n = 1132
0.0
97.7
86.4
**
embryonic survival [in %]
C
pcmd-1(syb975); mCherry::H2B
N2
PCMD-1 PBDm Nter
PCMD-1 PBDm Cter
PCMD-1 PDm
α-GFP
α-tubulin
D
cell cycle progression
n = 10
n = 9
ns
time from NEBD in P0 until NEBD in AB [sec]

**Figure S1**

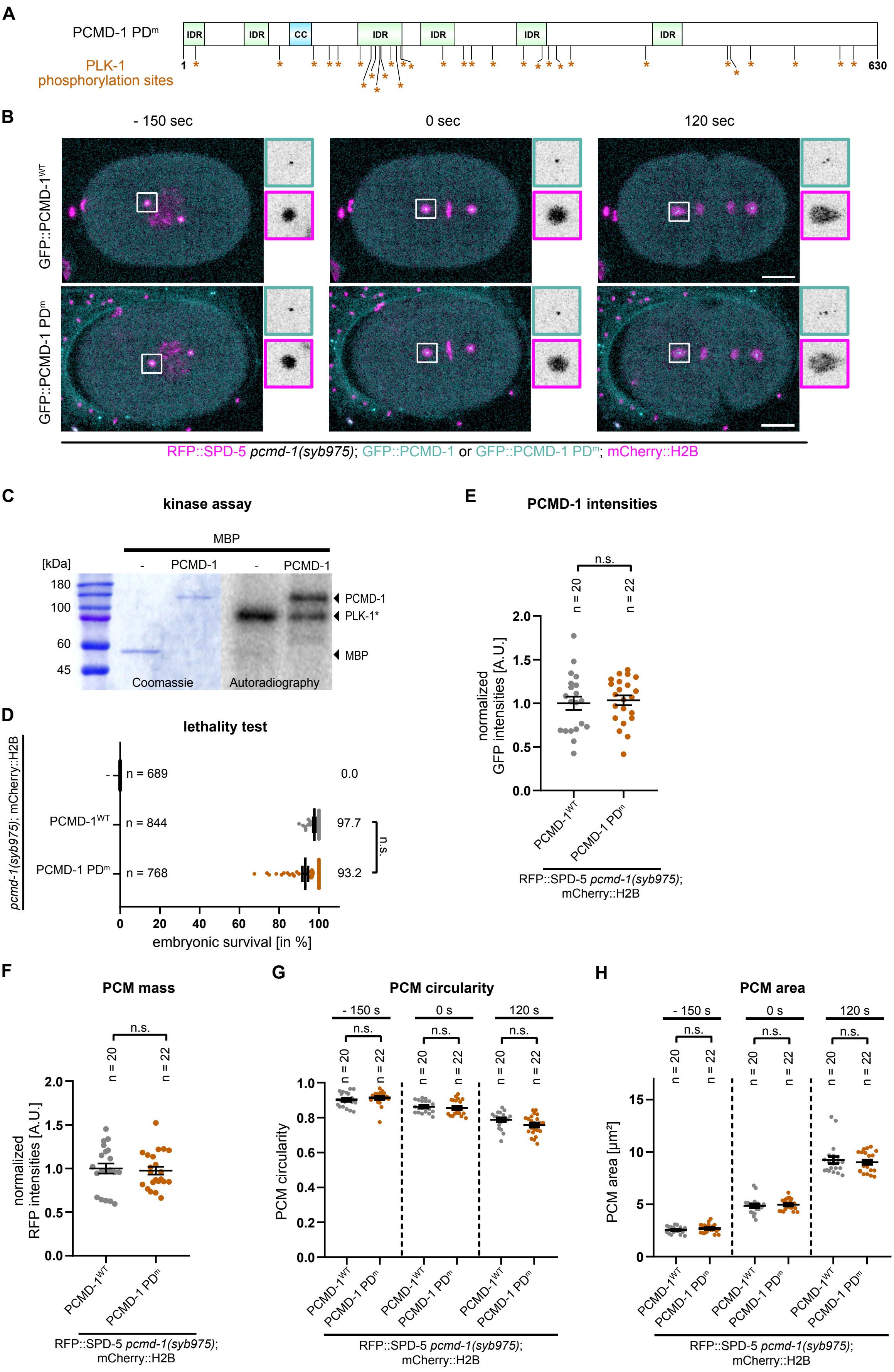
A
PCMD-1 PDm
PLK-1 phosphorylation sites
IDR
CC
1
630
B
- 150 sec
0 sec
120 sec
GFP::PCMD-1WT
GFP::PCMD-1 PDm
RFP::SPD-5 pcmd-1(syb975); GFP::PCMD-1 or GFP::PCMD-1 PDm; mCherry::H2B
C
kinase assay
MBP
[kDa]
-
PCMD-1
180
100
60
45
PCMD-1
PLK-1*
MBP
Coomassie
Autoradiography
D
lethality test
pcmd-1(syb975); mCherry::H2B
n = 689
0.0
PCMD-1WT
n = 844
97.7
PCMD-1 PDm
n = 768
93.2
n.s.
embryonic survival [in %]
E
PCMD-1 intensities
normalized GFP intensities [A.U.]
n = 20
n = 22
RFP::SPD-5 pcmd-1(syb975); mCherry::H2B
F
PCM mass
normalized RFP intensities [A.U.]
G
PCM circularity
- 150 s
0 s
120 s
H
PCM area
PCM area [µm²]

**Figure S2**

**A**

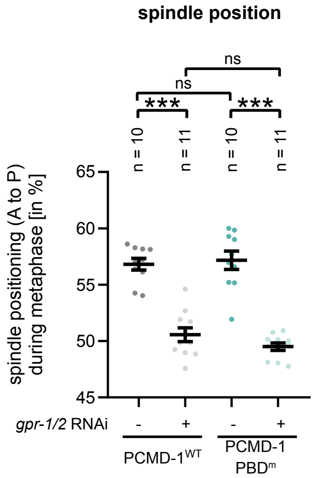
spindle position
ns
ns
***
***
n = 10
n = 11
n = 10
n = 11
spindle positioning (A to P) during metaphase [in %]
65
60
55
50
45
gpr-1/2 RNAi
-
+
-
+
PCMD-1WT
PCMD-1 PBDm

**Figure S3**

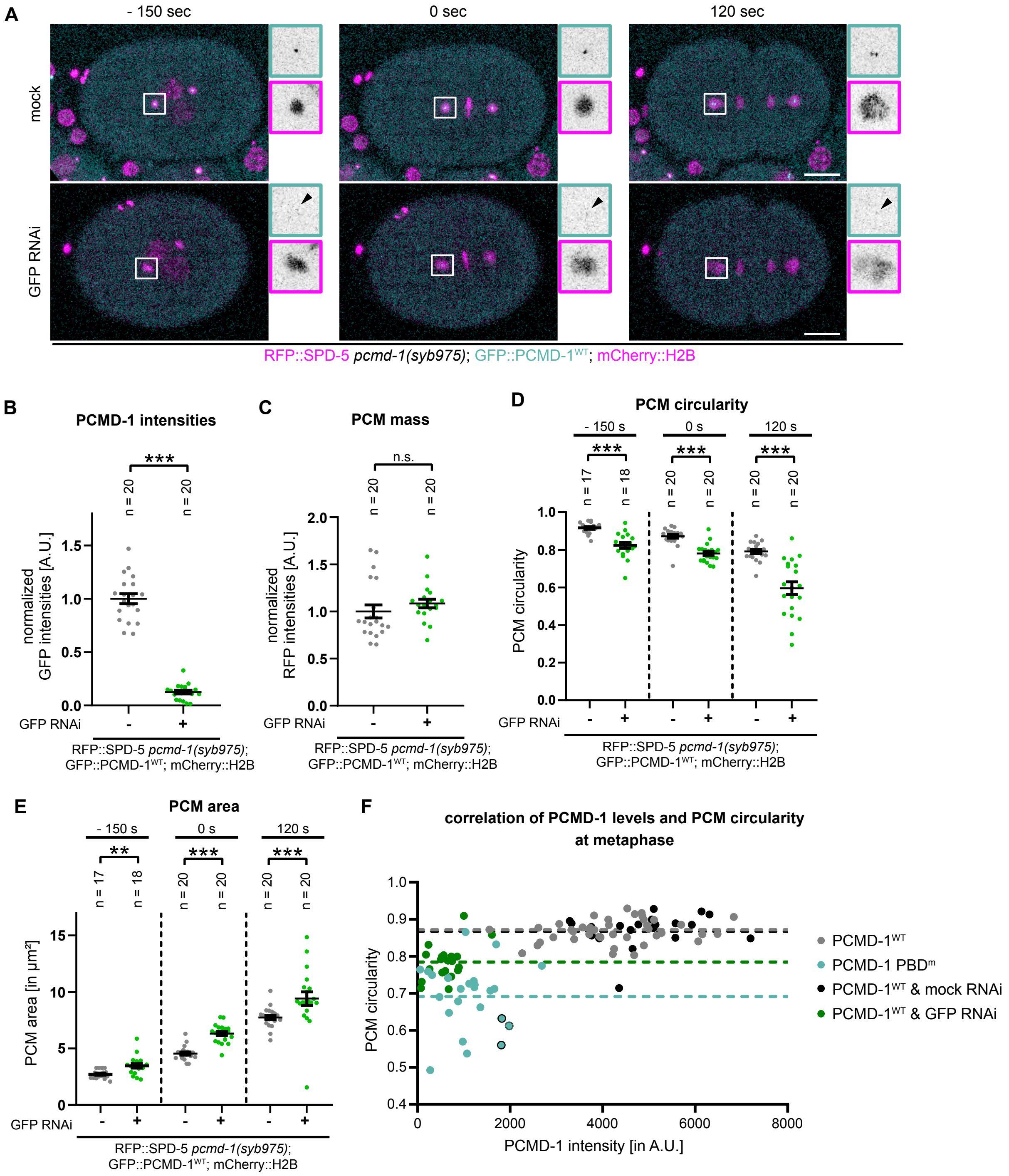
A
- 150 sec
0 sec
120 sec
mock
GFP RNAi
RFP::SPD-5 pcmd-1(syb975); GFP::PCMD-1WT; mCherry::H2B
B
PCMD-1 intensities
***
n = 20
n = 20
normalized GFP intensities [A.U.]
GFP RNAi
RFP::SPD-5 pcmd-1(syb975); GFP::PCMD-1WT; mCherry::H2B
C
PCM mass
n.s.
normalized RFP intensities [A.U.]
D
PCM circularity
- 150 s
0 s
120 s
n = 17
n = 18
PCM circularity
E
PCM area
**
PCM area [in µm²]
F
correlation of PCMD-1 levels and PCM circularity at metaphase
PCMD-1 intensity [in A.U.]
PCMD-1WT
PCMD-1 PBDm
PCMD-1WT & mock RNAi
PCMD-1WT & GFP RNAi

**Figure S4**

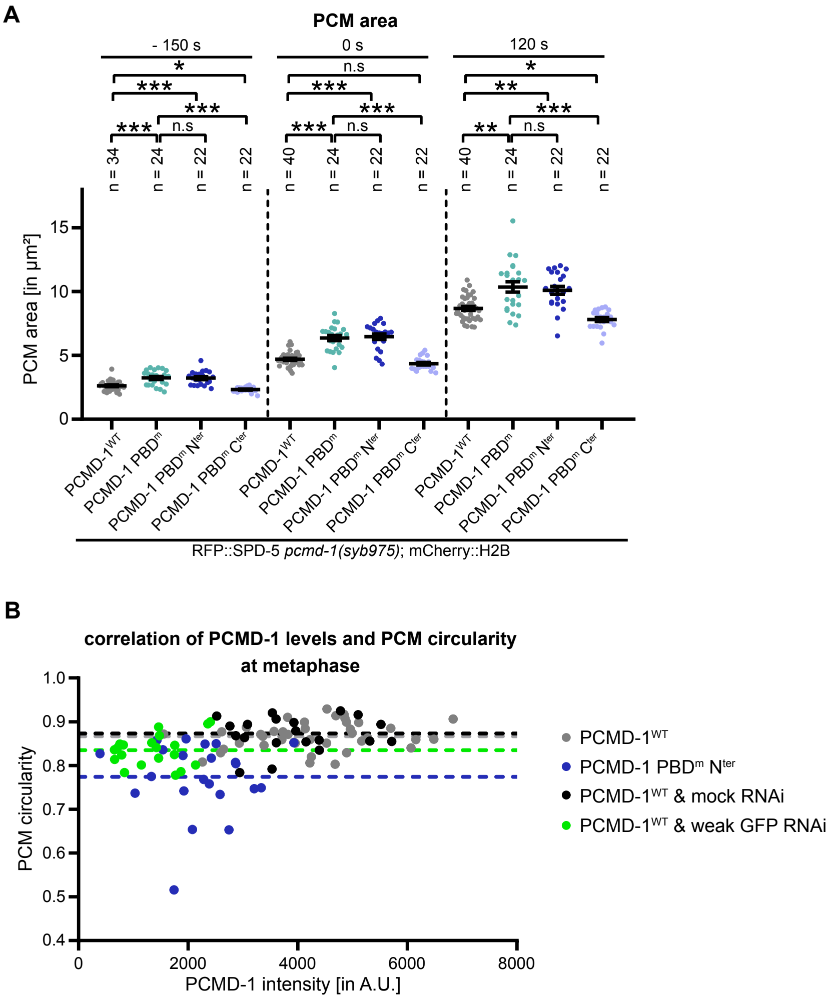
A
PCM area
- 150 s
0 s
120 s
PCM area [in µm²]
PCMD-1WT
PCMD-1 PBDm
PCMD-1 PBDm Nter
PCMD-1 PBDm Cter
RFP::SPD-5 pcmd-1(syb975); mCherry::H2B
B
correlation of PCMD-1 levels and PCM circularity at metaphase
PCM circularity
PCMD-1 intensity [in A.U.]
PCMD-1WT
PCMD-1 PBDm Nter
PCMD-1WT & mock RNAi
PCMD-1WT & weak GFP RNAi

**Figure S5**

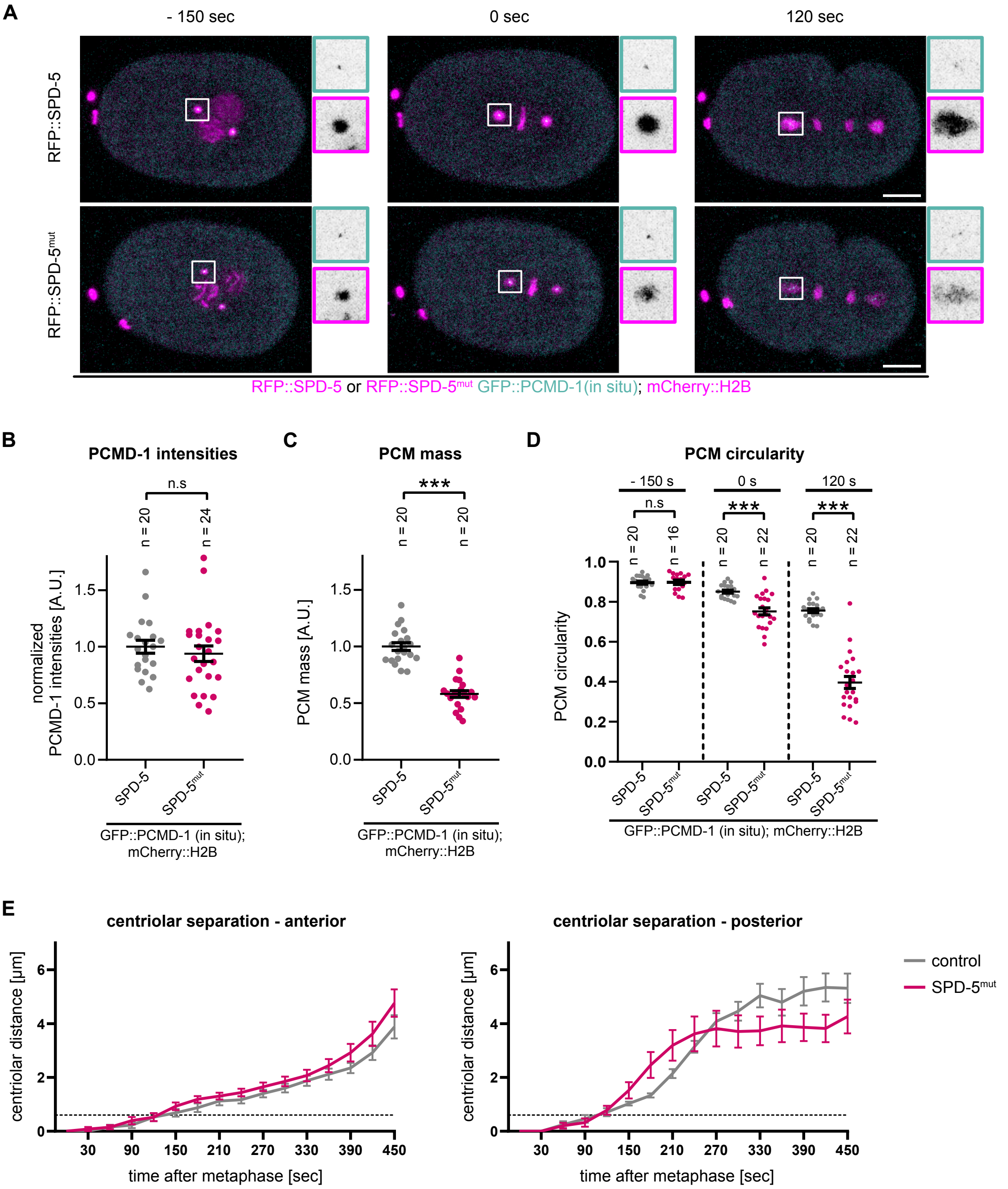

A
- 150 sec
0 sec
120 sec
RFP::SPD-5
RFP::SPD-5mut
RFP::SPD-5 or RFP::SPD-5mut GFP::PCMD-1(in situ); mCherry::H2B
B
PCMD-1 intensities
n.s
n = 20
n = 24
normalized PCMD-1 intensities [A.U.]
SPD-5
SPD-5mut
GFP::PCMD-1 (in situ); mCherry::H2B
C
PCM mass
***
n = 20
n = 20
PCM mass [A.U.]
SPD-5
SPD-5mut
GFP::PCMD-1 (in situ); mCherry::H2B
D
PCM circularity
- 150 s
0 s
120 s
n.s
***
***
n = 20
n = 16
n = 20
n = 22
n = 20
n = 22
PCM circularity
SPD-5
SPD-5mut
GFP::PCMD-1 (in situ); mCherry::H2B
E
centriolar separation - anterior
centriolar separation - posterior
centriolar distance [µm]
time after metaphase [sec]
control
SPD-5mut


**Figure S6**

**Mutated PCMD-1 PBD binding sites and PCMD-1 phosphorylation sites**

| position | PCMD-1 PBDm | PCMD-1 PBDm Nter | PCMD-1 PBDm Cter | PCMD-1 PDm |
|---|---|---|---|---|
| S12A | x | x | | x |
| S13A | x | x | | |
| S56A | x | x | | |
| S71A | x | x | | |
| S88A | x | x | | x |
| S119A | | | | x |
| S133A | | | | x |
| S134A | x | x | | |
| S138A | x | x | | |
| S139A | x | x | | |
| S141A | | | | x |
| S142A | x | x | | |
| S161A | | | | x |
| S171A | | | | x |
| T175A | | | | x |
| S179A | | | | x |
| T180A | x | x | | x |
| S185A | x | x | | |
| S189A | | | | x |
| T194A | | | | x |
| S198A | | | | x |
| S199A | x | x | | x |
| S217A | x | x | | |
| S226A | x | x | | |
| S227A | x | x | | |
| T228A | x | x | | |
| T236A | | | | x |
| S248A | x | x | | |
| S255A | | | | x |
| S262A | | | | x |
| T281A | | | | x |
| S309A | | | | x |
| S326A | | | | x |
| S332A | | | | x |
| T339A | | | | x |
| S352A | | | | x |
| T355A | x | | x | |
| S392A | x | | x | |
| T420A | | | | x |
| T453A | x | | x | |
| S491A | x | | x | |
| S494A | | | | x |

| | | | | |
|---|---|---|---|---|
| S497A | | | | x |
| S512A | x | | x | |
| S515A | | | | x |
| T555A | | | | x |
| T596A | x | | x | x |
| T608A | | | | x |
| **total #** | **23** | **17** | **6** | **30** |

**Table 1**

| strain name | *genotype* | refrence |
|---|---|---|
| N2 | Bristol isolate | CGC |
| TMD272 | *pcmd-1(syb975) I; mikSi34[pmai-2:GFP::PCMD-1_PLK1_PDmutant] II, ltIs37 [(pAA64) pie-1p::mCherry::his-58 + unc-119(+)] IV* | this study |
| TMD295 | *spd-5(wow36[tagrfp-t^3xmyc::spd-5]) pcmd-1(syb975) I; mikSi34[pmai-2:GFP::PCMD-1_PLK1_PDmutant] II; ltIs37 [(pAA64) pie-1p::mCherry::his-58 + unc-119(+)] IV* | this study |
| TMD296 | *spd-5(wow36[tagrfp-t^3xmyc::spd-5]) pcmd-1(syb975) I; mikSi37[pmai-2:GFP::PCMD-1_PLK1_PBDmutant]II; ltIs37 [(pAA64) pie-1p::mCherry::his-58 + unc-119(+)] IV* | this study |
| TMD306 | *pcmd-1(syb975) I; mikSi37[pmai-2:GFP::PCMD-1_PLK1_PBDmutant]II; ltIs37 [(pAA64) pie-1p::mCherry::his-58 + unc-119(+)] IV* | this study |
| TMD322 | *spd-5(wow36[tagrfp-t^3xmyc::spd-5]) pcmd-1(syb975) I; mikSi6[pmai-2:GFP::PCMD-1]II; ltIs37 [(pAA64) pie-1p::mCherry::his-58 + unc-119(+)] IV* | this study |
| TMD327 | *pcmd-1(syb975) I; ltIs37 [(pAA64) pie-1p::mCherry::his-58 + unc-119(+)] IV* | this study |
| TMD342 | *pcmd-1(syb975) I; mikSi39[pmai-2:GFP::PCMD-1]II; ltIs37 [(pAA64) pie-1p::mCherry::his-58 + unc-119(+)] IV* | this study |
| TMD356 | *pcmd-1(syb975) I; mikSi47[pmai-2:GFP::PCMD-1_PBD_N-ter] II; ltIs37 [(pAA64) pie-1p::mCherry::his-58 + unc-119(+)] IV* | this study |
| TMD374 | *spd-5(wow36[tagrfp-t^3xmyc::spd-5]) pcmd-1(syb975) I; mikSi47[pmai-2:GFP::PCMD-1_PBD_N-ter] II; ltIs37 [(pAA64) pie-1p::mCherry::his-58 + unc-119(+)] IV* | this study |
| TMD381 | *spd-5(wow36[tagrfp-t^3xmyc::spd-5]) pcmd-1(syb975) I; mikSi57[pmai-2:GFP::PCMD-1_PBD_C-ter] II; ltIs37 [(pAA64) pie-1p::mCherry::his-58 + unc-119(+)] IV* | this study |
| TMD382 | *pcmd-1(syb975) I; mikSi57[pmai-2:GFP::PCMD-1_PBD_C-ter] II; ltIs37 [(pAA64) pie-1p::mCherry::his-58 + unc-119(+)] IV* | this study |
| TMD383 | *spd-5(wow36[tagrfp-t^3xmyc::spd-5]) pcmd-1(syb486[gfp::pcmd-1]) I; ltIs37 [(pAA64) pie-1p::mCherry::his-58 + unc-119(+)] IV* | this study |
| TMD384 | *spd-5(wow36syb5737[D734-918]) pcmd-1(syb486[gfp::pcmd-1]) I; ltIs37 [(pAA64) pie-1p::mCherry::his-58 + unc-119(+)] IV* | this study |
| TMD385 | *pcmd-1(syb975) I; mikSi6[pmai-2:GFP::PCMD-1]II; plk-1(syb2781)::mCherry III* | this study |
| TMD386 | *pcmd-1(syb975) I; mikSi37[pmai-2:GFP::PCMD-1_PLK1_PBDmutant]II; plk-1(syb2781)::mCherry III* | this study |

**Table 2**
